# Regulation of cAMP accumulation and activity by distinct phosphodiesterase subtypes in INS-1 cells and human pancreatic β-cells

**DOI:** 10.1101/593418

**Authors:** Evan P.S. Pratt, Kyle E. Harvey, Amy E. Salyer, Shiqi Tang, Gregory H. Hockerman

## Abstract

Pancreatic β-cells express multiple phosphodiesterase (PDE) subtypes, but the specific roles for each in β-cell function, particularly in humans, is not clear. We evaluated the cellular role of PDE1, PDE3, and PDE4 activity in the rat insulinoma cell line INS-1 and in primary human β-cells using subtype-selective PDE inhibitors. Using a genetically encoded, FRET-based cAMP sensor, we found that the PDE1 inhibitor 8MM-IBMX and the PDE4 inhibitor rolipram elevated cAMP levels above baseline in the absence and presence of 18 mM glucose in INS-1 cells. Inhibition of PDE1 or PDE4 potentiated glucose-stimulated insulin secretion in INS-1 cells. In contrast, the inhibition of PDE3 with cilostamide had little effect on cAMP levels or glucose-stimulated insulin secretion. PDE1 inhibition, but not PDE3 of PDE4 inhibition, reduced palmitate-induced caspase-3/7 activation, and enhanced CREB phosphorylation in INS-1 cells. In human β-cells, only PDE3 or PDE4 inhibition increased cAMP levels in 1.7 mM glucose, but PDE1, PDE3, or PDE4 inhibition potentiated cAMP levels in 16.7 mM glucose. Inhibition of PDE1 or PDE4 increased cAMP levels to a greater extent in 16.7 mM glucose than in 1.7 mM glucose in human β-cells. In contrast, elevation of cAMP levels by PDE3 inhibition was not different at these glucose concentrations. PDE1 inhibition also potentiated insulin secretion from human islets, suggesting that the role of PDE1 may be conserved between INS-1 cells and human pancreatic β-cells. Our results suggest that inhibition of PDE1 may be a useful strategy to potentiate glucose-stimulated insulin secretion, and to protect β-cells from the toxic effects of excess fatty acids.

## Introduction

Pancreatic β-cells secrete the blood glucose-lowering hormone insulin to maintain glucose homeostasis in the body (1). Pancreatic β-cell dysfunction and cell death underlies the development of type 2 diabetes (2). At the cellular level, glucose-stimulated insulin secretion (GSIS) is driven by Ca^2+^ influx through the L-type voltage-gated Ca^2+^ channels (L-VGCC) Ca_v_1.2 and Ca_v_1.3 (3), and release of Ca^2+^ from the endoplasmic reticulum (ER) (4). GSIS is further regulated by the second messenger 3’,5’-cyclic adenosine monophosphate (cAMP), which is generated by the enzyme adenylyl cyclase (AC) (5). In addition to enhancing GSIS, cAMP promotes pancreatic β-cell mass through increased replication (6) and decreased apoptosis (7). Both glucose (8-10) and incretin hormones (11), such as glucagon-like peptide-1 (GLP-1), are capable of stimulating cAMP production and subsequent activation of the cAMP effector proteins Protein Kinase A (PKA) and Exchange Protein Directly Activated by cAMP (Epac) (12). PKA and Epac regulate insulin secretion through proximal effects on the machinery involved in exocytosis at the plasma membrane (13-15) and distal effects on ER Ca^2+^ release channels (16, 17). cAMP signaling is compartmentalized to microdomains within the cell, including near sites of ER Ca^2+^ release, by phosphodiesterase enzymes (PDE), which degrade cAMP to 5’-AMP.

PDE1, PDE3, PDE4, and PDE8 are widely-regarded as the primary PDE subtypes responsible for regulating cytosolic cAMP levels and GSIS in rodent β-cell lines, and rodent and human islets (18). PDE1 is the only subtype that is regulated by Ca^2+^/Calmodulin (19, 20) and is predicted to serve a critical role in pancreatic β-cells where Ca^2+^ dynamics and signaling are prominent (21). Indeed, the subtype-selective PDE1 inhibitor 8-methoxymethyl-3-isobutyl-1-methylxanthine (8MM-IBMX) elevated resting and glucose-stimulated cAMP levels and enhanced GSIS in βTC3 cells (21), MIN6 cells, primary mouse β-cells (22) and intact islets from rat and human (23). Yet, PDE3 appears to be the primary PDE in mouse, rat, and human islets, and the pancreatic β-cell lines INS-1 and MIN6. PDE3 overexpression led to reduced glucose-stimulated cAMP production, impaired GSIS (24, 25) and glucose intolerance (26), whereas PDE3 inhibition (22, 23, 27) or PDE3 knockdown (28, 29) elevated resting and glucose-stimulated cAMP levels and GSIS (30). It has also been suggested that leptin and insulin-like growth factor 1 attenuate insulin secretion through PDE3B activation (31, 32). Subtype-selective inhibitors and knockdown of PDE4 or PDE8 also elevated resting cAMP levels in INS-1 cells (28, 30), βTC3 cells (21), and intact rat and human islets (23, 27), suggesting that PDE4 and PDE8 may too be required. PDE8 knockdown enhanced GSIS in INS-1 cells (28, 30) and MIN6 cells (22), which directly correlates with PDE8-mediated regulation of resting cAMP levels. PDE4 inhibition, on the other hand, enhanced GSIS in clonal pancreatic β-cell lines (28, 30) but not in primary islets (23, 27), revealing a disconnect between clonal and primary β-cells. Furthermore, PDE4 activity was elevated in MIN6 cells and primary mouse β-cells following stimulation with glucose, as compared with resting conditions (22). The paucity of information regarding PDE activity in human β-cells and the disparate results in various rodent models have hampered the development of selective PDE inhibition as a strategy for treatment of type 2 diabetes.

Using subtype-selective inhibitors of PDE1, PDE3, PDE4, and PDE8, we performed a side-by-side comparison of the role of each subtype in INS-1 cells and for the first time, in single human pancreatic β-cells. We used a state-of-the-art fluorescent sensor of cAMP to evaluate PDE-mediated regulation of cAMP under resting conditions and during glucose stimulation. The latter allowed us to directly compare our findings with the ability of each PDE inhibitor to potentiate GSIS. Furthermore, we revealed a novel Ca^2+^-dependent mechanism by which PDE inhibition can potentiate GSIS in INS-1 cells. Finally, we show that PDE inhibition not only enhanced GSIS but also promoted INS-1 cell survival in the face of stress.

## Materials and Methods

### Chemicals gand Reagents

D-glucose was purchased from Mallinckrodt Chemicals (Dublin, Ireland). IBMX (3-Isobutyl-8-(methoxymethyl)-1-methyl-1H-purine-2,6(3H,7H)-dione) was purchased from Sigma-Aldrich (St. Louis, MO). 8MM-IBMX (3-Isobutyl-8-(methoxymethyl)-1-methyl-1H-purine-2,6(3H,7H)-dione) was purchased from Santa Cruz Biotechnology (Dallas, TX). Cilostamide (N-cyclohexyl-N-methyl-4-(1,2-dihydro-2-oxo-6-quinolyloxy) butyramide), rolipram (4-[3-(cyclopentyloxy)-4-methoxyphenyl]-2-pyrrolidinone), PF-04671536 (5-Methyl-3-[[(2R)-4 (thiazolylmethyl)-2-morpholino]methyl]-3H-1,2,3-trazolol[4,5]pyrimidin-7-amine), and forskolin were purchased from Tocris Bioscience (Bristol, UK). PF-04957325 (3-[[(2R)-4-(1,3-thiazol-2-ylmethyl)morpholin-2-yl]methyl]-5-(trifluoromethyl)triazolo[4,5-d]pyrimidin-7-amine) was purchased from MedChemExpress (Monmouth Junction, NJ). Unless otherwise indicated, all other reagents were purchased from Sigma-Aldrich.

### INS-1 Cell Culture

INS-1 cells were cultured in complete RPMI-1640 media containing 11 mM glucose, 10 mM HEPES, 10% fetal bovine serum (FBS), 11 mg/mL sodium pyruvate, 10,000 units/mL penicillin, 10,000 µg/mL streptomycin, and 50 µM β-mercaptoethanol at 37°C and 5% CO_2_ (33). In some experiments, INS-1 cells were serum-deprived overnight using minimal RPMI-1640 media containing 2.5 mM glucose, 10 mM HEPES, 11 mg/mL sodium pyruvate, 10,000 units/mL penicillin, 10,000 µg/mL streptomycin and 50 µM β-mercaptoethanol supplemented with 0.1% fatty acid-free bovine serum albumin (BSA).

### Intracellular Ca^2+^ Assay

INS-1 cells were plated in 96-well black-walled plates (Corning Life Sciences, Corning, NY) and incubated for at least 24h at 37°C and 5% CO_2_. Cells were washed once with 200 µL Phosphate Buffered Saline (PBS) and incubated in 100 µL Krebs-Ringer HEPES Buffer (KRBH) containing 5 µM Fura2-AM (Molecular Probes, Eugene, OR) for 1h at room temperature. KRBH contained 134 mM NaCl, 3.5 mM KCl, 1.2 mM KH_2_PO_4_, 0.5 mM MgSO_4_, 1.5 mM CaCl_2_, 5 mM NaHCO_3_ and 10 mM HEPES and was supplemented with 0.05% fatty-acid free BSA (pH 7.4). Following this 1h incubation, KRBH containing Fura2-AM was removed, and the cells were washed once with 200 µL KRBH. Cells were pretreated for 30 min with 100 µL KRBH containing the working concentration of inhibitors or KRBH alone.

Intracellular Ca^2+^ assays were performed using a Synergy 4 Multi-Mode Plate Reader (BioTek Instruments, Winooski, VT). Fura2 fluorescence was measured using a 508/20 nm band-pass emission filter and 340/11 nm or 380/20 nm band-pass excitation filters. For acute application of caffeine or PDE inhibitors, a 15 sec baseline was recorded prior to injection of 100 µL of a 2X concentration of stimulant, and fluorescence was recorded for an additional 2 min.

Fluorescence was collected from both channels every 0.7 sec. For long-term stimulation with PDE Inhibitors in combination with glucose or glucose + forskolin, a 5 min baseline was recorded prior to injection of 100 µL of a 2X concentration of stimulant, and fluorescence was recorded for an additional 1h. Fluorescence was collected from both channels every 0.7 sec. To assess changes in cytosolic Ca^2+^, the fluorescent signal collected using the 340 nm excitation filter was divided by the fluorescent signal collected using the 380 nm excitation filter (340/380 nm) for each well over the entire time-course. The average ratio during the first min was subtracted from the entire time-lapse to yield a baseline of zero. Each treatment was performed in quadruplicate, and the average 340/380 nm ± SE was determined for the time-course. At least three independent experiments were performed for each treatment, and the average AUC ± SE was determined.

### Insulin Secretion

INS-1 cells were plated in 24-well plates (Corning Life Sciences, Corning, NY) and incubated for at least 24h at 37°C and 5% CO_2_. Cells were washed once with PBS and incubated with 500 µL KRBH alone or containing the working concentration of inhibitors for 30 min at 37°C and 5% CO_2_. Following pretreatment, 500 µL KRBH containing a 2X concentration of glucose or glucose + PDE inhibitors along with the working concentration of the same inhibitors used in the pretreatment was added to each well and incubated for an additional 1h at 37°C and 5% CO_2_. Following this incubation, the supernatant (1 mL) was transferred from each well to 1.5 mL Eppendorf tubes and stored at 20°C until assayed. Secreted insulin was measured using the High-Range Insulin ELISA kit (ALPCO, Salem, NH), according to the manufacturer’s instructions. After supernatant removal, cells from each well were immediately lysed in 200 µL Cell lysis buffer (20 mM Na_2_HPO_4_ + 150 mM NaCl + 0.1% Triton X-100, pH 7.4) + freshly added protease inhibitors (800 nM aprotinin, 50 µM leupeptin, 1 µg/mL pepstatin, 1 mM benzamidine, 1 mM 4-(2-Aminoethyl) benzenesulfonylfluoride,10 µg/mL calpain inhibitor I and 10 µg/mL calpain inhibitor II). Cell lysis buffer was transferred from each well to 1.5 mL tubes, placed on ice for 20 min and centrifuged at 14,000 × g for 10 min. The supernatant was transferred to fresh 1.5 mL tubes and stored at −20°C until assayed. Cellular protein was measured using the Pierce BCA Protein Assay Kit (Thermo Fisher Scientific, Waltham, MA), according to the manufacturer’s instructions. Each sample was assayed in duplicate. Secreted insulin was normalized to protein content, both of which were measured using the PowerWave Plate Reader (BioTek Instruments, Winooski, VT). A calibration curve was generated using standards provided in each kit, and the amount of secreted insulin and protein content were determined. The amount of insulin secreted (ng) was divided by protein content (mg) to yield an insulin/protein ratio for each well. Each treatment was performed in triplicate, and the average insulin/protein ratio was calculated among like wells. The average insulin/protein ratio of each treatment was normalized to the untreated condition to yield percent basal. This value was again normalized to the glucose treatment to yield percent glucose. At least three independent experiments were performed for each treatment, and the average percent glucose ± SE was determined.

### CREB Phosphorylation Assay

INS-1 cells were plated in 96-well black-walled plates and incubated for at least 24h at 37°C and 5% CO_2_. Cells were incubated for an additional 24h in minimal RPMI media. Cells were washed twice with PBS and incubated for 2h in KRBH at 37°C and 5% CO_2_. The preincubation buffer was removed and replaced with KRBH alone or containing the working concentration of inhibitors for 30 min at 37°C and 5% CO_2_. Cells were treated for 10 min with KRBH alone or KRBH containing the working concentration of PDE inhibitors. Following this incubation, cells were fixed with 4% formaldehyde and stored at 4°C until assayed. Total CREB and pCREB were measured using the Human/Mouse/Rat Phospho-CREB (S133) Cell-based ELISA (R&D Systems, Minneapolis, MN), according to the manufacturer’s instructions. Using a Synergy 4 Multi-Mode Plate Reader (BioTek Instruments, Winooski, VT), total CREB was measured at 450 nm with excitation at 360 nm, and pCREB was measured at 600 nm with excitation at 540 nm. The fluorescent signal of pCREB was divided by the fluorescent signal of total CREB to yield a pCREB/total CREB ratio for each well. Each treatment was performed in duplicate, and the average pCREB/total CREB was calculated between these two wells. The average pCREB/total CREB among background wells (no primary antibody) was subtracted from the values obtained from other wells. The average pCREB/total CREB ratio of each treatment was normalized to the untreated condition to yield percent basal. In some cases, the basal value was subtracted and the resulting values were normalized to the glucose treatment to yield percent glucose. At least three independent experiments were performed for each treatment, and the average percent basal or percent glucose ± SE was calculated.

### Caspase-3/7 Glo Assay

INS-1 cells were plated in 96-well white-walled plates (Corning Life Sciences, Corning, NY) and incubated for at least 24h at 37°C and 5% CO_2_. INS-1 cells were treated with a 3:1 molar ratio of palmitate to BSA, with and without the working concentration of PDE inhibitors, or BSA alone (untreated control). To prepare the stock solution of palmitate, sodium palmitate (100 mM) (Santa Cruz Biotechnology, Dallas, TX) was dissolved in 50% ethanol by incubating at 55°C for 10-15 min with frequent vortexing. As an untreated control, 50% ethanol that did not contain palmitate was also prepared. A 1:200 dilution (50 µL) of palmitate (500 µM final) or control solution (0.25% ethanol final) was added to 10 mL minimal RPMI media containing 1% fatty acid-free BSA. Tubes containing palmitate + BSA or BSA alone were incubated in a 37°C water bath for 30 min to promote conjugation. INS-1 cells were treated for 12h at 37°C and 5% CO_2_. Following incubation, treatments were removed and replaced with 50 µL of fresh media. 50 µL of Caspase-3/7 Glo Reagent (ProMega, Madison, WI) was added to each well to yield a 100 µL total volume. As a background control, 50 µL RPMI-1640 + 50 µL Caspase-3/7 Glo Reagent were added to several wells that did not contain INS-1 cells. The plate was shaken at 500 rpm for 30 sec and incubated in the dark at room temperature for 2h. Luminescence was recorded using a Synergy 4 Multi-Mode Plate Reader (BioTek Instruments, Winooski, VT). A 1s integration time was used in all experiments; however, the sensitivity was changed to achieve an average luminescence recording of 500 in the background wells. Each treatment was performed in triplicate, and the average luminescence signal was calculated among these wells. The average luminescent signal of each treatment was normalized to the untreated condition to yield percent basal. This value was again normalized to palmitate to yield percent palmitate. At least three independent experiments were performed for each treatment, and the average percent palmitate ± SE was determined.

### Human pancreatic islets

Human islets were acquired from the Integrated Islet Distribution Program (City of Hope National Medical Center, Duarte, CA). The average purity and viability of the human islet preparations used in these studies was 92% and 94%, respectively. All the donors were classified as nondiabetics, and the cause of death was reported as stroke (4 donors), head trauma (4 donors), anoxia (2 donors), or CNS tumor (1 donor). The average age was 40 years old (range 28-62), the average weight was 225.3 pounds (range 170-296) and the average body mass index was 33.0 (range 23.7-49.9). Upon receipt, human islets were centrifuged at 200 × g for 5 min and re-suspended in RPMI-1640 media containing 11 mM glucose, 10 mM HEPES, 10% FBS, 10,000 units/mL penicillin and 10,000 µg/mL streptomycin (human islet media). Islets were plated in 24-well plates (30-40 islets/well) for FRET-based imaging experiments or ultra-low attachment 60 mm culture dishes (Corning Life Sciences, Corning, NY) for single islet insulin secretion assay. Islets were incubated for at least 24h at 37°C and 5% CO_2_ for recovery.

### Dissociation and transfection of human pancreatic islets

Human islets were dissociated from one well of a 24-well plate (30-40 islets). Islets were transferred to a 1.5 mL Eppendorf tube and centrifuged at 200 × g for 5 min. The media was carefully removed and replaced with 1 mL warm Versene (Thermo Fisher Scientific, Waltham, MA). The islets were re-suspended by pipetting up and down 5X, and centrifuged at 200 × g for 5 min. The Versene was carefully removed and replaced with 1 mL warm Accutase (Thermo Fisher Scientific, Waltham, MA). The islets were re-suspended by pipetting up and down 5X and placed on a rocker at room temperature for 5 min for constant agitation. Following agitation, the islets were briefly placed in a 37°C water bath and re-suspended by pipetting up and down 5X. The 1.5 mL tube was placed on the rocker at room temperature for an additional 5 min. Following agitation, the islets were briefly warmed in a 37°C water bath and re-suspended for a final time by pipetting up and down 5X. The dissociated islets were pelleted at 200 × g for 5 min, and the Accutase solution was carefully removed and replaced with 100 µL human islet media. The islets were resuspended and transferred to the center of a 40 mm coverslip coated with poly-D-lysine placed in a 60 mm dish. The dissociated islet cells were allowed to settle for 10-15 min prior to transfection. 500 ng Epac-S^H187^ + 1 µL P3000 + 1 µL Lipofectamine 3000 (Life Technologies, Grand Island, NY) was used for transfection. The DNA + P3000 and lipofectamine 3000 were added to separate 1.5 mL tubes, each containing 10 µL OPTI-MEM. The contents of both tubes were combined to yield a total volume of 20 µL, and the DNA and lipofectamine were allowed to complex at room temperature for 5 min. Following 4-6h incubation at 37°C and 5% CO_2_, fresh human islet media was carefully added to the 60 mm dish as not to disturb the dissociated islets. Dissociated cells were imaged at 48h post-transfection.

### Single human islet insulin secretion assay

Using a dissection microscope, single human islets (50-100 µm in diameter) were transferred from an ultra-low attachment 60 mm dish to 96-well V-bottom clear-walled plate (Corning Life Sciences, Corning, NY) containing 100 µL human islet media and incubated for 48h at 37°C and 5% CO_2_. Following this incubation, the cells were washed once with 50 µL warm KRBH + 1.7 mM glucose and incubated with 100 µL warm KRBH + 1.7 mM glucose for 30 min at 37°C and 5% CO_2_. The pretreatment was removed and replaced with 100 µL warm KRBH + 1.7 mM glucose (1.7G) or KRBH + 16.7 mM glucose (16.7G) with and without the working concentration of PDE inhibitors, and incubated for an additional 1h at 37°C and 5% CO_2_. The supernatants containing secreted insulin were transferred to fresh wells on the same 96-well V-bottom plate. Single human islets were lysed using 100 µL RIPA Buffer (50 mM Tris + 150 mM NaCl + 1% Triton X-100 + 0.5% Sodium Deoxycholate + 0.1% SDS) + freshly added protease inhibitors. The 96-well V-bottom plate containing cell lysate and secreted insulin was wrapped in parafilm and stored at 20°C until assayed. Secreted insulin and insulin content were diluted 1:10 and 1:100, respectively, to fall in the dynamic range of the kit, and each sample was assayed in duplicate using the Human Insulin Chemiluminescence ELISA (ALPCO, Salem, NH), according to the manufacturer’s instructions. Within 5-15 min of adding the chemiluminescent substrate, luminescence was detected using the Synergy 4 Multi-Mode Plate Reader using a 1s integration time and detector sensitivity of 150. A calibration curve was generated using standards provided in the kit, and the amount of secreted insulin and insulin content were determined. The resulting values were corrected for dilution, and the amount of secreted insulin was divided by total insulin (secreted insulin + insulin content) to yield a secreted insulin/total insulin percentage for each well. Each treatment was performed in triplicate, and the average % secreted/total insulin ± SE was calculated.

### FRET-based cAMP imaging

To measure cAMP accumulation in the presence of PDE inhibitors, INS-1 cells were transfected with 1 µg Epac-S^H187^ DNA using 2.5 µL Lipofectamine 2000 (Life Technologies, Grand Island, NY), according to the manufacturer’s instructions. Following 4-6h incubation at 37°C and 5% CO_2_, the cells were split into 60 mm dishes containing 40 mm glass coverslips coated with poly-D-lysine and imaged at 48h post-transfection. INS-1 cells and dissociated human islet cells were imaged using a Nikon A1 Confocal and Perfect Focus Ti-E Inverted Microscope equipped with an Apo TIRF 60x Oil DIC N2 (NA 1.49) objective lens. 40 mm coverslips were assembled into a RC-31 imaging chamber (Warner instruments, Hamden, CT) and attached to a six-channel perfusion mini-valve system (VC-6M). The donor signal of Epac-S^H187^ was collected using the following confocal parameters: 457 nm laser line of a Multi-Argon laser (457/476/488/514 nm), a 400-457/514 nm primary dichroic mirror, a 520 nm long-pass dichroic mirror and a 485/35 nm band-pass emission filter. The FRET signal of Epac-S^H187^ was collected using the following confocal parameters: 457 nm laser line of a Multi-Argon laser (457/476/488/514 nm), a 400-457/514 nm primary dichroic mirror, a 565 nm long-pass dichroic mirror and a 538/33 nm band-pass emission filter. Confocal images of donor and FRET signal were collected sequentially every 2 sec using the following parameters: 512 x 512 total pixels, 4.8 µs pixel dwell, pinhole size of 1.2 AU and varied zoom factor, laser power and detector gain. A baseline signal was collected for 1 min prior to stimulation. KRBH containing the working concentration of IBMX or subtype-selective PDE inhibitors were perfused over the cells, and once a new baseline was achieved, KRBH was used as a wash. For experiments with INS-1 cells, KRBH in the basal or “unstimulated” condition contained no glucose. For experiments with human β-cells, basal KRBH contained 1.7 mM glucose. Time-lapse recordings of donor and FRET signal were analyzed using NIS Elements 4.0 software. To assess changes in cAMP accumulation, the ratio of donor to FRET signal of Epac-S^H187^ was determined for each cell over the time-course, to yield several single cell traces. Each trace was normalized to the average ratio during the first min, to yield a baseline of one. At least three independent experiments were performed for each treatment. The percent change from baseline was determined for each treatment and the average ± SE was determined.

### Data Analysis

Data were analyzed using GraphPad Prism 7. Data are represented as the average ± SE. Statistical analysis was performed using one-way or two-way ANOVA with the Tukey post-hoc test. Alternatively, the student’s unpaired t-test was used. *P* < 0.05 was considered significant.

## Results

### PDE-mediated regulation of resting and glucose-stimulated cAMP levels in INS-1 cells

The role of PDE1, PDE3, PDE4 and PDE8 in regulating both resting and glucose-stimulated cAMP levels in pancreatic β-cells is poorly understood. We used the Epac1-based, high-affinity, high-dynamic range FRET sensor Epac-S^H187^ to evaluate the effect of subtype-selective PDE inhibitors on cAMP levels in INS-1 cells with real-time resolution (34). We transiently transfected INS-1 cells with Epac-S^H187^ and imaged cells by confocal microscopy at 48 hours post-transfection in KRBH without glucose. Application of the pan PDE inhibitor IBMX (100 µM) resulted in a robust increase in cAMP levels (34.99 ± 1.94%) that recovered to baseline upon drug removal (Fig 1A). We sequentially applied the subtype-selective inhibitors 8MM-IBMX, cilostamide, and rolipram to test the role of the IBMX-sensitive PDE1, PDE3, and PDE4, respectively (Fig 1A). The concentrations of PDE inhibitors used in this study were 5-20 times the reported IC_50_ values (35-37) and were comparable with those used in a similar study performed in MIN6 cells (22). Application of 100 µM 8MM-IBMX (17.87 ± 1.42%) and 10 µM rolipram (10.6 ± 2.14%) significantly elevated cAMP levels above baseline, whereas 1 µM cilostamide (5.04 ± 0.5%) had little effect (Fig 1B). Notably, 8MM-IBMX had a significantly greater effect on resting cAMP levels than either cilostamide and rolipram. Furthermore, the rolipram-induced increase in cAMP levels was significantly greater than cilostamide. Thus, PDE1, PDE3, PDE4 regulate resting cAMP levels in INS-1 cells with a rank order of PDE1 > PDE4 > PDE3. Summation of the percent increase in cAMP levels elicited by each subtype-selective PDE inhibitor was 33.5%, which is comparable to the increase observed with IBMX. Since PDE8 is not inhibited by IBMX (38), we examined the effect of the PDE8-selective inhibitor PF-04957325 (39) on resting cAMP levels in INS-1 cells. We found that 100 nM PF-04957325 (IC50 < 1 nM for both PDE8A and 8B) had no effect on cAMP levels under basal conditions (S1 Fig). Additionally, the PDE8-selective inhibitor PF-04671536 (IC_50_ < 2 nM for both PDE8A and 8B)(42), didn’t increase basal cAMP levels in INS-1 cells at concentrations up to 5 μM (S1 Fig).

**Figure 1.**
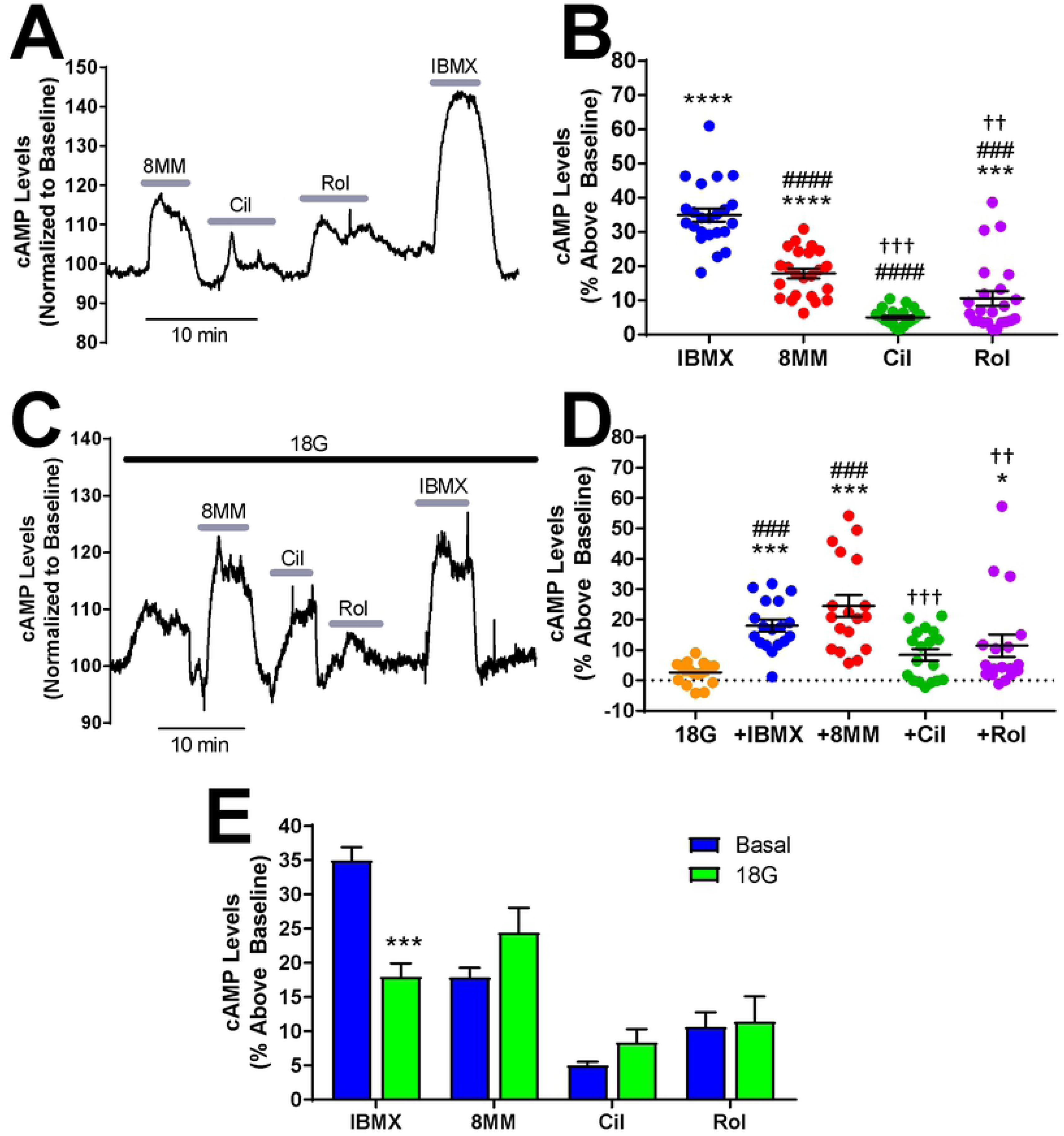
PDE1 and PDE4 regulate basal cAMP levels and glucose-stimulated cAMP in INS-1 cells. **A)** Subtype-selective PDE inhibitors and the pan PDE inhibitor IBMX raise basal cAMP levels in INS-1 cells. The Epac-S^H187^ mTurquoise2/FRET ratio is directly proportional to the cAMP level. Data shown is a representative experiment from a single cell. **B)** The percent increase in cAMP levels above baseline elicited by PDE inhibitors in INS-1 cells under basal conditions. IBMX (100 µM), 8MM-IBMX (100 µM) and rolipram (10 µM) significantly elevate cAMP levels above baseline. Further, the percent increase elicited by each of the subtype-selective inhibitors is significantly less than that of IBMX. Last, 8MM-IBMX caused a significantly greater elevation in cAMP levels than either cilostamide (1 µM) or rolipram (***, P < 0.001 compared to baseline; ###, P < 0.001 compared to IBMX; †††, P < 0.001, ††, P < 0.01 compared to 8MM-IBMX; One-way ANOVA, Tukey post-hoc test). Data shown are mean ± SE from 23 cells taken from four independent experiments. **C)** Subtype-selective PDE inhibitors and the pan PDE inhibitor IBMX raise cAMP levels in the presence of 18 mM glucose in INS-1 cells. Data shown is a representative experiment from a single cell. **D)** The percent increase in cAMP levels above baseline elicited by PDE inhibitors in INS-1 cells in the presence of 18 mM glucose. IBMX, 8MM-IBMX and rolipram significantly elevate cAMP levels above baseline, whereas only IBMX and 8MM-IBMX are significantly greater than glucose. Last, 8MM-IBMX caused a significantly greater elevation in cAMP levels than either cilostamide or rolipram. Data shown are mean ± SE from 18 cells taken from four independent experiments. (***, P < 0.001, *, P < 0.05 compared to baseline; ###, P < 0.001 compared to 18G; †††, P < 0.001, ††, P < 0.01 compared to 8MM-IBMX; One-way ANOVA, Tukey post-hoc test). **E)** Comparison of cAMP accumulation stimulated by PDE inhibitors in the absence and presence of 18 mM glucose. IBMX stimulated a greater increase in cAMP levels in the absence of glucose (basal), compared to 18 mM glucose (***, *P* < 0.001; Student’s unpaired t-test).

PDE1 activity is upregulated by Ca^2+^/Calmodulin (19, 20), while PDE3 and PDE4 activities are upregulated by PKA-mediated phosphorylation (40). Glucose elevates Ca^2+^ and cAMP levels in pancreatic β-cells (8); therefore, it’s plausible that the PDE profile is altered in INS-1 cells following glucose stimulation. To test this, we stimulated INS-1 cells expressing Epac-S^H187^ with 18 mM glucose and sequentially applied the PDE inhibitors. Fig 1C shows initial application of glucose (18 mM) resulted in a small but detectable rise in cytosolic cAMP levels (2.64 ± 0.83%). Similar to our findings under resting conditions, 100 µM IBMX (18 ± 1.93%), 100 µM 8MM-IBMX (24.45 ± 3.61%), and 10 µM rolipram (11.43 ± 3.66%) significantly elevated cAMP levels in the presence of glucose, while 1 µM cilostamide (8.39 ± 1.92%) did not (Fig 1D). Furthermore, 8MM-IBMX had a significantly greater effect on glucose-stimulated cAMP levels compared with rolipram and cilostamide, suggesting that PDE1 is the primary regulator of cAMP levels under resting and glucose-stimulated conditions. In addition, we directly compared the effect of each PDE inhibitor under resting conditions with glucose stimulation. We found that the PDE1-selective inhibitor 8MM-IBMX and PDE3-selective inhibitor cilostamide trended toward increased cAMP levels in the presence of glucose, but this difference did not reach statistical significance (Fig 1E). Interestingly, the IBMX-induced rise in cAMP levels was significantly greater in the absence of glucose than in 18 mM glucose.

### PDE-mediated regulation of resting and glucose-stimulated cAMP levels in human β-cells

Given that the rat-derived INS-1 cell line may not reflect human β-cell physiology, we examined the roles of PDE1, PDE3, PDE4 and PDE8 in regulation of resting cAMP levels in isolated pancreatic β-cells dissociated from human islets. Previous reports have demonstrated that PDE1, PDE3 and PDE4 regulate cAMP levels in intact human islets; however, there is no evidence which suggests that this is specific to β-cells (23, 41). Dissociated human islet cells were transiently transfected with Epac-S^H187^ and imaged at 48 hours post-transfection by confocal microscopy. To distinguish pancreatic β-cells from other islet cell types, we employed a previously established pharmacological approach (22). Unlike other islet cell types, pancreatic β-cells express both the GLP-1 receptor and α2-adrenergic receptor, which when activated with their respective agonists, lead to AC activation and inhibition, respectively. At the end of each imaging experiment, we applied the GLP-1R agonist GLP-1 (50 nM) followed by co-administration of the α_2_-adrenergic receptor agonist clonidine (1 µM). If a cell exhibited a GLP-1-induced rise in cAMP levels that was dampened or abolished by application of clonidine, it was classified as a pancreatic β-cell (Fig 2A-C). We used this approach to measure cAMP levels in human pancreatic β-cells obtained from non-diabetic human donors (S1 Table) and found that, as in INS-1 cells, application of 100 µM IBMX in low glucose (1.7 mM) resulted in a robust increase in cAMP levels (29.45 ± 3.17%) (Fig 2). Further, 1 µM cilostamide (15.3 ± 2.94%) and 10 µM rolipram (12.53 ± 4.43%) significantly elevated resting cAMP levels, whereas 100 µM 8MM-IBMX (9.35 ± 1.53%) did not (Fig 2). This suggests that unlike INS-1 cells, PDE3 and PDE4, but not PDE1, regulates resting cAMP levels in human pancreatic β-cells. We also examined the role of PDE8 in regulating cAMP levels in human pancreatic β-cells. We used the PDE8 inhibitor PF-04671536 since it is reported to stimulate GSIS in human islets (42). However, 5 μM PF-04671536 had no effect on basal cAMP levels in human pancreatic β-cells (S2. Fig).

**Figure 2.**
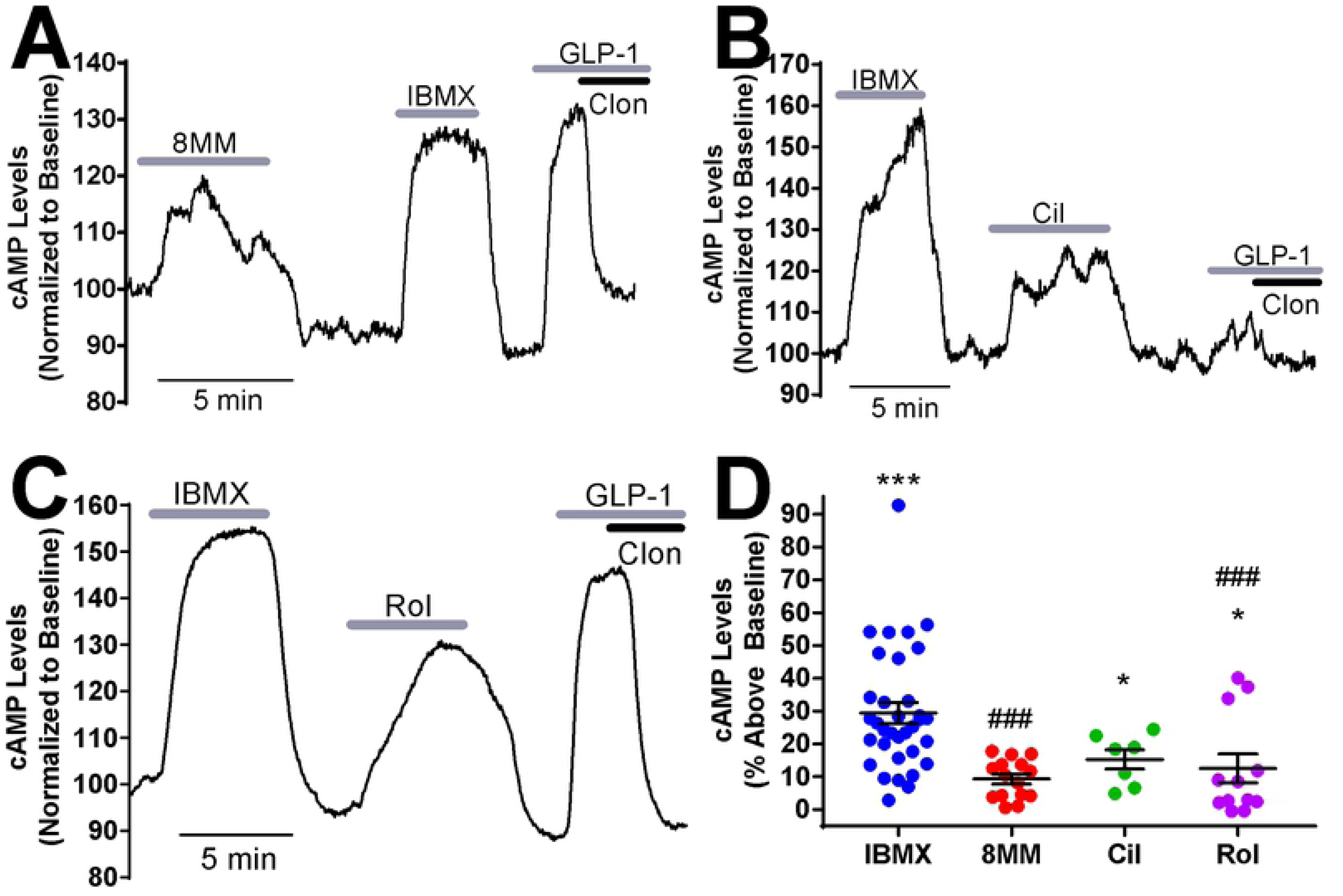
Regulation of basal cAMP levels by PDEs in human pancreatic β-cells. **A-C)** Subtype-selective PDE inhibitors and IBMX raise cAMP levels in pancreatic β-cells dissociated from human islets under basal conditions (1.7 mM glucose). For each experiment, IBMX (100 µM) and either 8MM-IBMX (100 µM) (**A**), cilostamide (1 µM) (**B**) or rolipram (10 µM) (**C**) were perfused onto the cell. Pancreatic β-cells were identified at the end of each experiment by addition of GLP-1 (50 nM) with and without the α2-receptor agonist clonidine (1 µM). Data shown are representative experiments from single human pancreatic β-cells. **D)** The percent increase in cAMP levels above baseline elicited by PDE inhibitors in human pancreatic β-cells under basal conditions. IBMX, cilostamide and rolipram significantly elevate cAMP levels above baseline. The percent increase in cAMP levels stimulated by 8MM-IBMX and rolipram is significantly less than that of IBMX (***, P < 0.001, *, P < 0.05 compared to baseline; ###, P < 0.001 compared to IBMX; One-way ANOVA, Tukey post-hoc test). Data shown are average ± SE from 15 cells (8MM-IBMX), 7 cells (cilostamide), 12 cells (rolipram) and 34 cells (IBMX), collected from four human islet preparations.

To compare regulation of resting and glucose-stimulated cAMP levels by PDEs in human pancreatic β-cells, we examined the role of PDE1, PDE3, and PDE4 in regulation of cAMP levels during 16.7 mM glucose stimulation in human pancreatic β-cells from non-diabetic human donors (S1 Table). At the end of each experiment, pancreatic β-cells were again identified by the characteristic increase in cAMP concentration stimulated by GLP-1 which is rapidly reversed by clonidine (Fig 3A-C). We found that the pan PDE inhibitor IBMX (48.08 ± 3.25%), PDE1-selective inhibitor 8MM-IBMX (25.81 ± 2.75%), PDE3-selective inhibitor cilostamide (15.99 ± 1.4%), and PDE4-selective inhibitor rolipram (27.58 ± 5.77%) each significantly elevated cAMP levels in the presence of glucose (Fig 3D). A direct comparison of PDE-mediated regulation of cAMP levels under resting (Fig 2) and stimulatory conditions (Fig 3) revealed that the IBMX-induced rise was significantly elevated in the presence of glucose (Fig 3E). Consistent with this, 8MM-IBMX and rolipram had a significantly greater effect in the presence of glucose compared with resting conditions, whereas cilostamide did not. Thus, PDE1 and PDE4 are upregulated following glucose stimulation, suggesting that the activity of these PDE subtypes is enhanced in the presence of glucose. In the case of PDE1, activity is highly dependent on glucose stimulation in human pancreatic β-cells.

**Figure 3.**
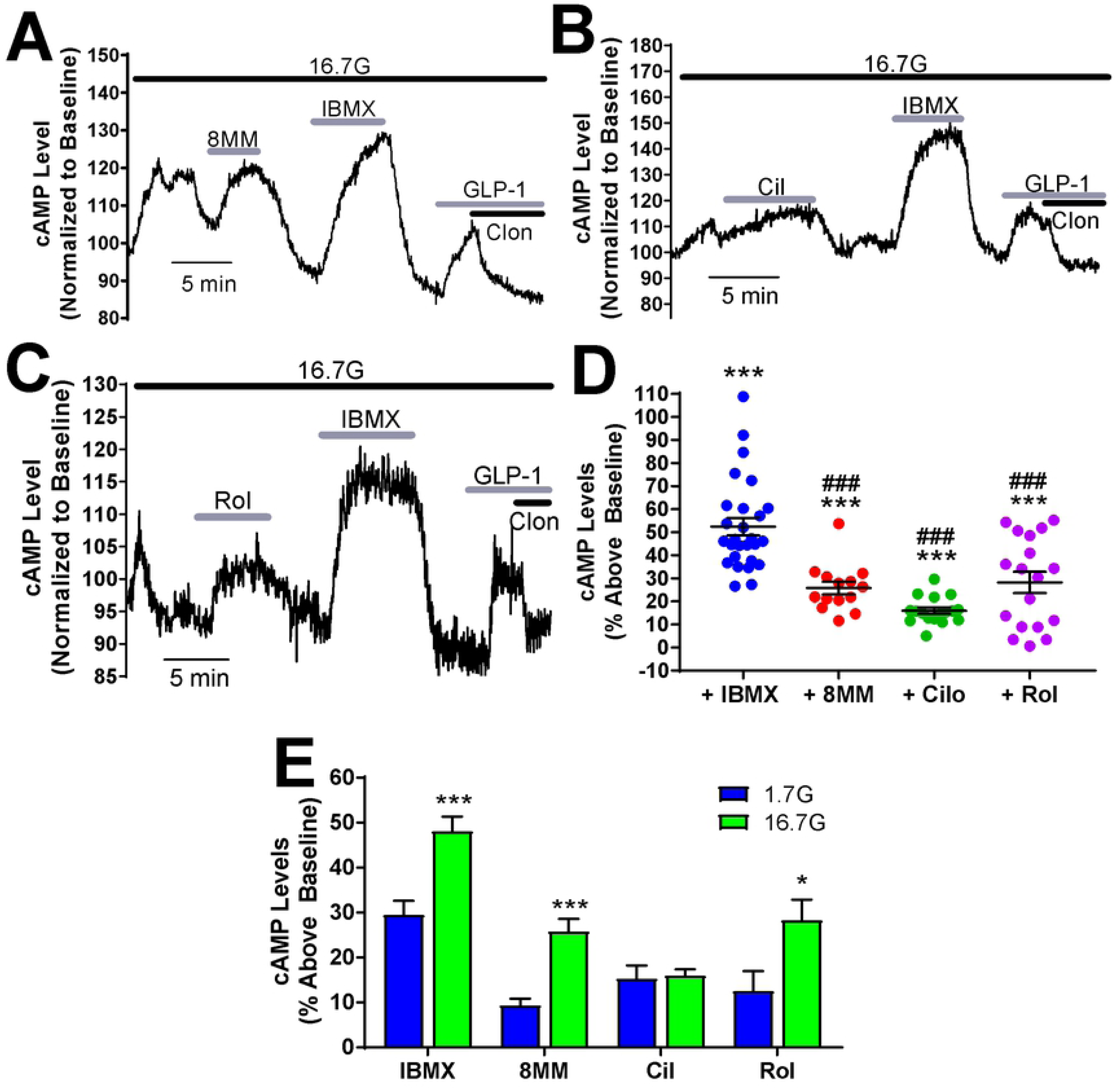
Regulation of glucose-stimulated cAMP by PDEs in human pancreatic β-cells. **A-C)** Subtype-selective PDE inhibitors and IBMX raise cAMP levels in pancreatic β-cells dissociated from human islets under high glucose (16.7 mM) stimulation conditions. For each experiment, IBMX (100 µM) and either 8MM-IBMX (100 µM) (**A**), cilostamide (1 µM) (**B**) or rolipram (10 µM) (**C**) were perfused onto the cell. Pancreatic β-cells were identified using GLP-1 and clonidine as described for Figure 5. Data shown are representative experiments from single human pancreatic β-cells. **D)** The percent increase in cAMP levels above baseline elicited by PDE inhibitors in human pancreatic β-cells in the presence of 16.7 mM glucose. IBMX, 8MM-IBMX, cilostamide, and rolipram all significantly elevate cAMP levels above baseline (***, *P* < 0.001; ^###^, *P* < 0.001 compared to IBMX; One-way ANOVA with Tukey post-hoc test). Data are shown as mean ± SE from 27 cells (IBMX), 14 cells (8MM-IBMX), 17 cells (cilostamide), and 18 cells (rolipram). **E)** Comparison of cAMP accumulation stimulated by PDE inhibitors in 1.7 mM (basal) or 18 Mm glucose. IBMX, 8MM-IBMX, and rolipram stimulated a greater increase in cAMP levels in 18 mM glucose, compared to 1.7 mM glucose (***, *P* < 0.001, *, *P* < 0.01; Student’s unpaired t-test).

### PDE-mediated regulation of GSIS in INS-1 cells and human islets

Elevated cAMP concentrations can enhance GSIS (43); therefore, we tested whether PDE-mediated regulation of glucose-stimulated cAMP levels in INS-1 cells (Fig 1) correlates with the ability of subtype-selective PDE inhibitors to potentiate GSIS. Here, we measured insulin secretion from INS-1 cells stimulated with 18 mM glucose in the absence or presence of PDE inhibitors. 18 mM glucose significantly stimulated insulin secretion from INS-1 cells compared to basal secretion (Fig 4A). Each experiment was normalized to insulin secretion in the presence of 18 mM glucose alone. Treatment with 100 µM IBMX resulted in greater than a three-fold increase (3.24 ± 0.17) in insulin secretion in INS-1 cells compared with glucose alone (Fig 4B). Treatment of INS-1 cells with 18 mM glucose + 100 µM 8MM-IBMX (2.25 ± 0.15) or 10 µM rolipram (1.79 ± 0.1) significantly potentiated insulin secretion compared with glucose alone, whereas 1 µM cilostamide (1.12 ± 0.13) did not (Fig 4B). Moreover, potentiation of GSIS by cilostamide was significantly less than that observed with IBMX, 8MM-IBMX, or rolipram. This suggests that PDE1 and PDE4, but not PDE3, regulate GSIS in INS-1 cells, consistent with our measurements of cAMP stimulation in Fig 1D, showing that PDE1 is the major subtype degrading cAMP in the presence of 18 mM glucose, followed closely by PDE4. Lastly, we found that PDE8-selective inhibitor PF-04957325 (100 nM) did not significantly potentiate GSIS in INS-1 cells (S1 Fig). Thus, it appears that the major IBMX-insensitive PDE isoform found in pancreatic β-cells does not play a significant role in either regulation of basal cAMP levels or GSIS in INS-1 cells.

**Figure 4.**
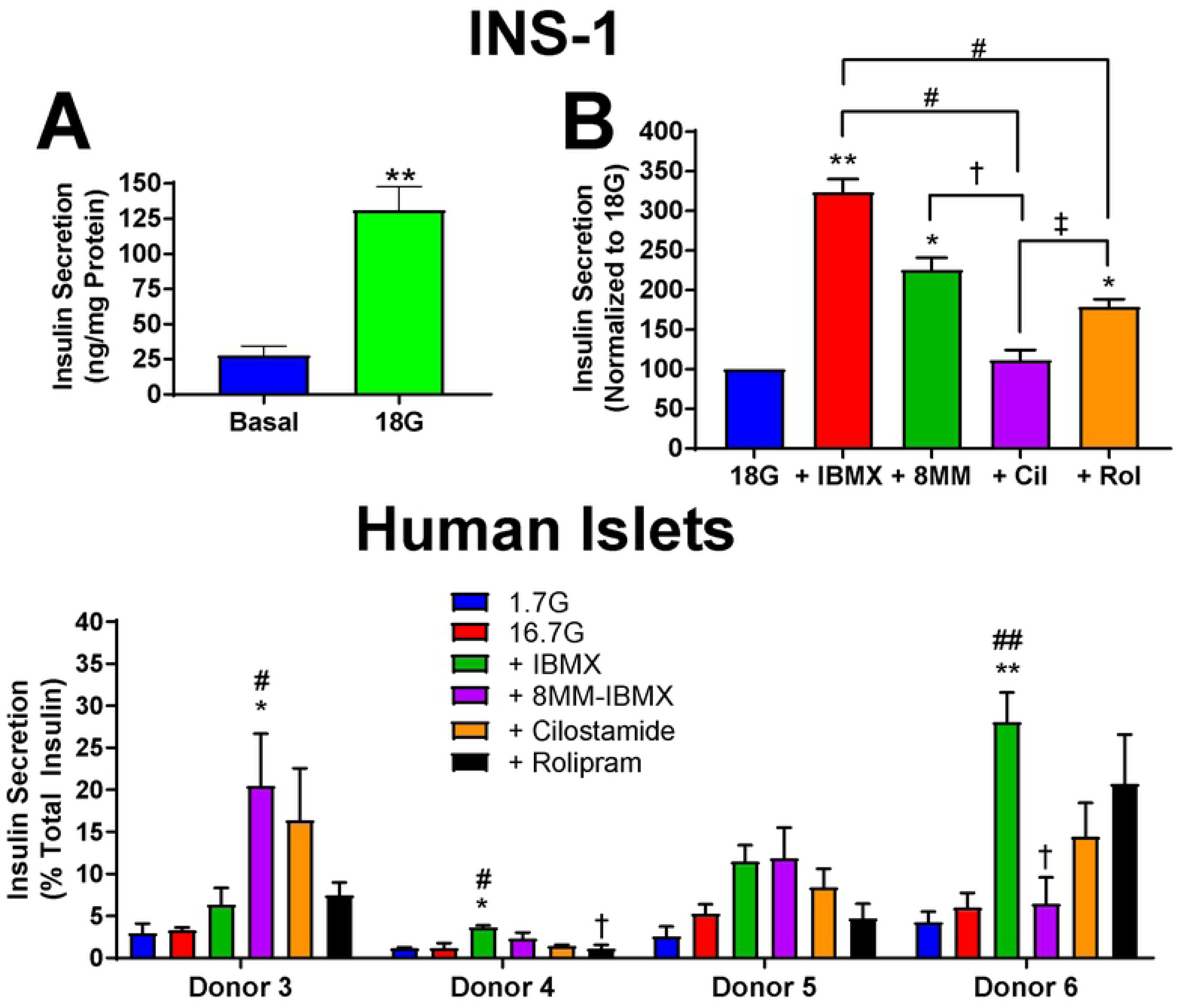
Effect of PDE inhibitors on GSIS in INS-1 cells and human pancreatic islets. **A)** Glucose (18 mM) significantly stimulates insulin secretion above basal levels in INS-1 cells (**, P < 0.01, compared to 0 glucose; Student’s unpaired t-test). Data shown are mean ± SE from four independent experiments. **B)** Subtype-selective PDE inhibitors and pan the PDE inhibitor IBMX potentiate insulin secretion stimulated with glucose (18 mM) in INS-1 cells. Each experiment was normalized to the glucose response. IBMX (100 µM), 8MM-IBMX (100 µM) and rolipram (10 µM) significantly stimulate insulin secretion, compared with glucose alone. Potentiation of GSIS with cilostamide (1 µM) or rolipram is significantly less than IBMX. Further, insulin secretion stimulated with cilostamide is significantly less than both 8MM-IBMX and rolipram (**, P < 0.01, *, P < 0.05 compared to 18G; #, P < 0.05 compared to IBMX; †, P < 0.05 compared to 8MM-IBMX; ‡, P < 0.05 compared to rolipram; One-way ANOVA, Tukey post-hoc test). Data shown are mean ± SE from four independent experiments. **C)** Subtype-selective PDE inhibitors and the pan PDE inhibitor IBMX potentiate insulin secretion stimulated with 16.7G in intact human islets isolated from four separate donors. In donor 3, 8MM-IBMX was significantly greater than 1.7G and 16.7G. In donor 4, IBMX was significantly greater than 1.7G and 16.7G, whereas rolipram was significantly less than IBMX. There are no significant differences among the treatment conditions for donor 5. Finally, in donor 6 we observed that IBMX had a significant effect above 1.7G and 16.7G, and 8MM-IBMX was less than IBMX (**, P < 0.01, *, P < 0.05 compared to 1.7G; ##, P < 0.01, #, P < 0.05 compared to 16.7G; †, P < 0.05 compared to IBMX; One-way ANOVA, Tukey post-hoc test). Data shown are mean ± SE from four human islet donors, in which each treatment was performed in triplicate.

Given the distinct effects of PDE subtype inhibition on cAMP levels in human pancreatic β-cells, we evaluated whether inhibitors of PDE1, PDE3, and PDE4 potentiate GSIS in isolated human islets using a single-islet insulin secretion assay (44). In short, islets obtained from non-diabetic human donors (S1 Table) were plated in a 96-well V-bottom plate, so that each well held a single islet. Human islets were incubated with PDE inhibitors at low glucose (1.7 mM) for 30 min prior to a 1h incubation with high glucose (16.7 mM) with and without PDE inhibitors. Insulin release was normalized to the insulin content of each islet. We found that 16.7 mM glucose stimulated a subtle increase in insulin secretion compared with 1.7 mM glucose in all four donors (Fig 4C), ranging from a 1.11-fold increase to 2.03-fold (S1 Table). However, in none of the donors did this apparent increase reach statistical significance. In contrast, the PDE inhibitors had more robust effects on GSIS in some donors. The pan PDE inhibitor IBMX trended toward potentiated GSIS in all four donors, and this reached statistical significance in two of the four donors (donors 4 and 6). The PDE1-selective inhibitor 8MM-IBMX enhanced GSIS in three donors (donors 3, 4 and 5), and this increase reached statistical significance in donor 3. The PDE3-selective inhibitor cilostamide trended toward potentiation of GSIS in three (donors 3, 5 and 6) and the PDE4-selective inhibitor rolipram trended toward significance in two (donors 3 and 6), but none of these increases reached statistical significance. Taken together, PDE1 appears to play a role in regulating GSIS in intact human islets, though there is clearly heterogeneity amongst individuals.

### PDE-mediated regulation of Ca^2+^ dynamics in INS-1 cells

8MM-IBMX and rolipram potentiated GSIS in INS-1 cells (Fig 4B), suggesting that PDE1 and PDE4 are the major PDE subtypes that regulate GSIS in INS-1 cells. PDE-mediated regulation of GSIS presumably occurs following the rise in cytosolic cAMP levels (Fig 1) and subsequent activation of the cAMP effector proteins, PKA (45) and Epac (46). However, both IBMX and 8MM-IBMX are derivatives of xanthine, as is caffeine, a potent activator of the ryanodine receptor (RyR) (47). Therefore, we examined whether or not, like caffeine, IBMX and 8MM-IBMX, and potentially cilostamide and rolipram, enhanced GSIS by stimulating RyR-mediated Ca^2+^ release. Using the fluorescent Ca^2+^ sensor Fura2-acetoxymethyl (AM), we found that acute application of caffeine (5 mM) stimulated a rapid and robust increase in cytosolic Ca^2+^ (Fig 5A&B). In contrast, application of IBMX and each of the subtype-selective PDE inhibitors, at the concentrations used to stimulate cAMP accumulation, did not markedly elevate cytosolic Ca^2+^ levels in INS-1 cells. Quantification of the area under the curve (AUC) showed that all of the of the PDE inhibitors stimulated a significantly smaller increase in intracellular Ca^2+^ concentration compared to caffeine: 100 μM IBMX (14.86 ± 4.9%), 100 μM 8MM-IBMX (23.9 ± 2.23%), 1 μM cilostamide (22.67 ± 1.92%) or 10 μM rolipram (15.22 ± 2.41%). Thus, potentiation of GSIS by IBMX, 8MM-IBMX and rolipram in INS-1 cells is likely not due to direct activation of RyR. Alternatively, it’s possible that PDE-mediated enhancement of GSIS occurs via activation of PKA and Epac, which can enhance ER Ca^2+^ release (16, 17). As expected, 18 mM glucose stimulated a rise in cytosolic Ca^2+^ that peaked approximately 10 min post-stimulation (Fig 5C). However, co-application of IBMX or any of the subtype-selective PDE inhibitors did not further increase intracellular Ca^2+^ levels in comparison with glucose alone (Fig 5D). Since the increase in cAMP levels stimulated by glucose alone is relatively modest, it’s possible that a more intense stimulation of cAMP levels might potentiate the Ca^2+^ response to glucose. Therefore, in an attempt to amplify the changes in cytosolic Ca^2+^, we stimulated INS-1 cells with glucose (18 mM) + the AC activator forskolin (25 µM). Glucose + forskolin stimulated a biphasic Ca^2+^ transient in INS-1 cells characterized by a rapid early peak followed by a prolonged plateau phase (Fig 5E). Co-application of the pan PDE inhibitor IBMX (150 ± 27.5%) and PDE1-selective inhibitor 8MM-IBMX (162 ± 15.48%) selectively enhanced the first phase of the Ca^2+^ transient stimulated with glucose + forskolin (Fig 5E&F). Notably, we’ve previously demonstrated the first phase of the tolbutamide-stimulated Ca^2+^ transient is due to ER Ca^2+^ release (48). In contrast, neither the PDE3-selective inhibitor cilostamide (110 ± 6.99%) nor the PDE4-selective inhibitor rolipram (123 ± 10.2%) had any effect. Thus, under conditions that strongly elevate Ca^2+^ and cAMP levels, PDE1 inhibition strongly enhances intracellular Ca^2+^ levels, providing a potential Ca^2+^-dependent mechanism by which PDE1 potentiates GSIS.

**Figure 5.**
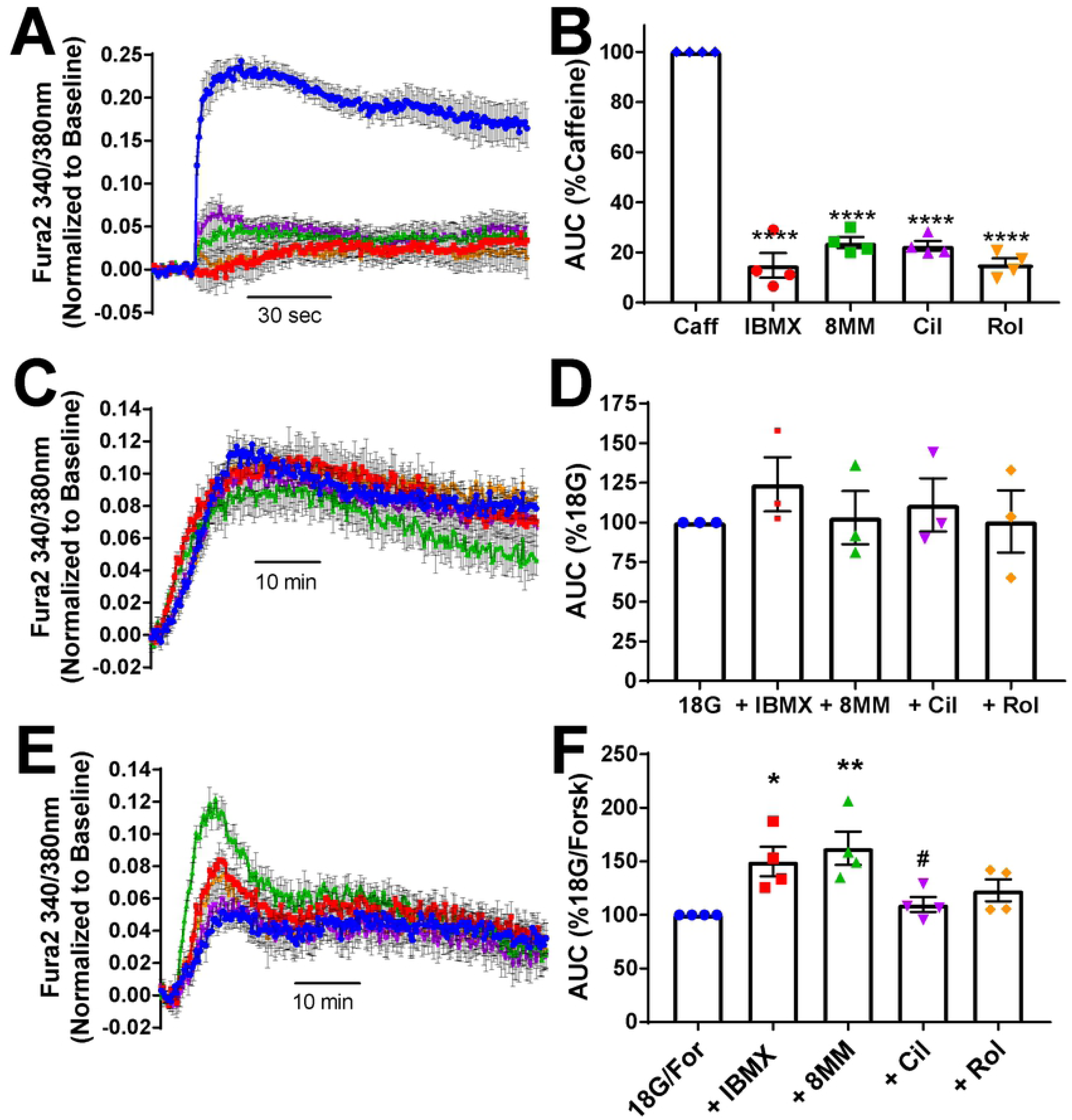
Effect of PDE inhibitors on intracellular Ca^2+^ dynamics in INS-1 cells. **A)** Caffeine robustly elevates intracellular Ca^2+^ in INS-1 cells when added acutely; however, the subtype-selective PDE inhibitors and the pan PDE inhibitor IBMX have little effect. Data shown are the mean ± SE from representative experiments performed in quadruplicate. **B)** AUC analysis of Ca^2+^ dynamics stimulated with caffeine (5 mM), 8MM-IBMX (100 µM), cilostamide (1 µM), rolipram (10 µM) or IBMX (100 µM). Each experiment was normalized to the response elicited by caffeine. The Ca^2+^ response elicited by each of the subtype-selective PDE inhibitors as well as IBMX was significantly less than that of caffeine (****, P < 0.0001 compared to caffeine; One-way ANOVA, Tukey post-hoc test). Data shown are average ± SE from four independent experiments. **C)** Glucose (18 mM) elevates intracellular Ca^2+^ levels in INS-1 cells but the subtype-selective PDE inhibitors and the pan PDE inhibitor IBMX have no additional effect. Data shown are the mean ± SE from representative experiments performed in quadruplicate. **D)** AUC analysis of Ca^2+^ dynamics stimulated with glucose (18 mM) alone and in the presence of either 8MM-IBMX (100 µM), cilostamide (1 µM), rolipram (10 µM) or IBMX (100 µM). Each experiment was normalized to the response elicited by 18G. There was no significant difference among the treatment conditions. Data shown are average ± SE from four independent experiments. **E)** Glucose (18 mM) + forskolin (25 µM) strongly elevates intracellular Ca^2+^ levels in INS-1 cells. The PDE1 inhibitor 8MM-IBMX (100 µM) and IBMX (100 µM) potentiate the initial phase of the response. Data shown are the mean ± SE from representative experiments performed in quadruplicate. **F)** AUC analysis of Ca^2+^ dynamics stimulated with glucose (18 mM) + forskolin (25 µM) alone and in the presence of either 8MM-IBMX (100 µM), cilostamide (1 µM), rolipram (10 µM) or IBMX (100 µM). Each experiment was normalized to the response elicited by 18G + forskolin. Addition of IBMX or 8MM-IBMX elicited a significantly greater elevation in intracellular Ca^2+^ (AUC) than 18G + forskolin alone. Ca^2+^ AUC stimulated by 18G+forskolin in the presence of cilostamide is significantly less than that in the presence of 8MM-IBMX (**, P < 0.01, *, P < 0.05 compared to 18G + forskolin; #, P < 0.05 compared to 8MM-IBMX; One-way ANOVA, Tukey post-hoc test). Data shown are average ± SE from four independent experiments.

### PDE-mediated regulation of INS-1 cell survival

cAMP not only enhances insulin secretion from pancreatic β-cells but also is an important regulator of β-cell mass, enhancing proliferation and cell survival (49). Not surprisingly, cAMP signaling is impaired in diabetic pancreatic β-cells (50). Given the robust effects of PDE inhibitors on cAMP levels (Fig 1), we tested whether raising cAMP levels using subtype-selective PDE inhibitors can rescue INS-1 cells from apoptosis caused by the saturated fatty acid palmitate, which is a widely-used model of lipotoxicity associated with type 2 diabetes (51). We exposed INS-1 cells to 500 µM palmitate at a 3:1 molar ratio with 1% BSA (52) in the presence of PDE inhibitors and measured caspase-3/7 activation, an early marker of apoptosis, at 12 hours post-treatment (53). Palmitate treatment significantly upregulated caspase-3/7 activity, compared with BSA alone (Fig 6A). Co-treatment of INS-1 cells with 100 µM IBMX (61.35 ± 8.65%) significantly decreased palmitate-induced caspase-3/7 activation (Fig 6B). Furthermore, we found that 100 µM 8MM-IBMX (64.81 ± 7.94%), but neither 1 µM cilostamide (87.54 ± 3.31%) nor 10 µM rolipram (75.45 ± 9.84%), significantly decreased palmitate-induced caspase 3/7 activation.

**Figure 6.**
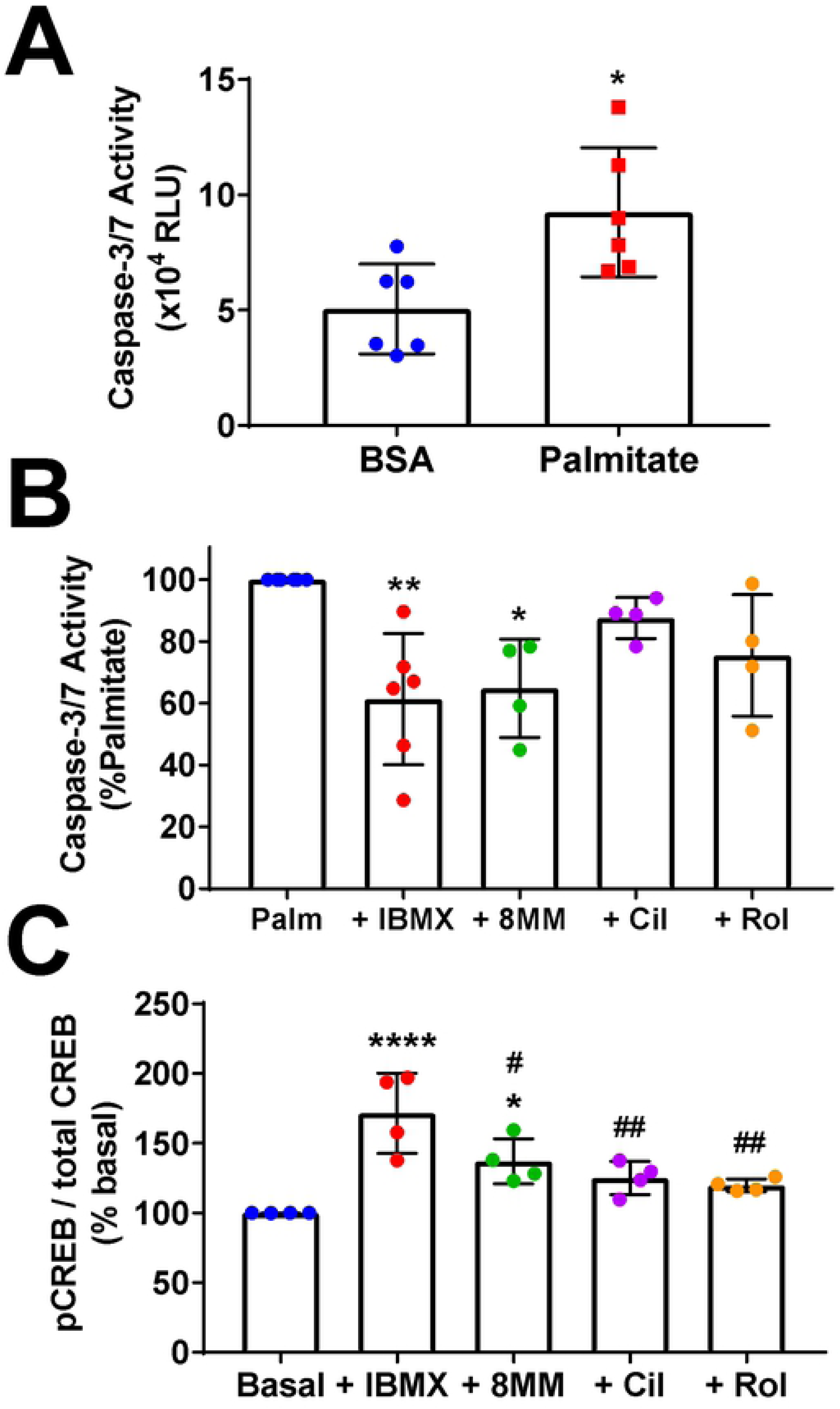
PDE1 inhibition reduces palmitate-induced apoptosis and stimulates CREB phosphorylation in INS-1 cells-. **A)** Incubation of INS-1 cells with palmitate for 12h significantly elevates caspase-3/7 activation (*, P < 0.05 compared to BSA; Student’s unpaired t-test). Data shown are average ± SE from six independent experiments. **B)** The PDE1 inhibitor 8MM-IBMX (100 µM) and pan PDE inhibitor IBMX (100 µM) significantly decrease the level of caspase-3/7 activation induced by palmitate toxicity (**, P < 0.01, *, P < 0.05 compared to palmitate; One-way ANOVA, Tukey post-hoc test). Each experiment was normalized to the level of palmitate toxicity. Data shown are average ± SE from 4-6 independent experiments. **C)** The PDE1 inhibitor 8MM-IBMX (100 µM) and pan PDE inhibitor IBMX (100 µM) significantly stimulate CREB phosphorylation above basal levels (****, P < 0.0001, *, P < 0.05 compared to basal; ##, P < 0.01, #, P < 0.05 compared to IBMX; One-way ANOVA, Tukey post-hoc test). Data shown are average ± SE from four independent experiments.

cAMP-response element binding protein (CREB) is a cAMP-dependent transcription factor that regulates gene transcription and pancreatic β-cell survival following PKA-dependent phosphorylation (7). Thus, a potential mechanism by which IBMX and 8MM-IBMX may activate pro-survival pathways in INS-1 cells is through an elevation in cAMP levels (Fig 1) that ultimately leads to PKA-mediated phosphorylation of CREB. To determine whether PDE inhibition stimulates CREB phosphorylation, we treated INS-1 cells with IBMX and the subtype-selective PDE inhibitors for 10 min, lysed the cells and measured CREB phosphorylation at S133 (pCREB) and total CREB expression using a commercially available ELISA kit. We calculated the ratio of pCREB to total CREB and normalized each experiment to basal pCREB/total CREB. Using this approach, we found that 100 µM IBMX (171 ± 14.32%) and 100 µM 8MM-IBMX (137 ± 8.05%) significantly elevated pCREB/total CREB, compared with basal conditions (Fig 6C). In contrast, neither 1 µM cilostamide (125 ± 5.94% basal) nor 10 µM rolipram (120 ± 2.27% basal) had a significant effect. Taken together, the pro-survival effect of the PDE-selective inhibitor 8MM-IBMX could be the result of elevated cAMP levels (Fig 1) and subsequent activation of the cAMP-responsive transcription factor CREB.

## Discussion

In this study, we examined the role of PDE1, PDE3, PDE4, and PDE8 in regulating resting and glucose-stimulated cAMP levels and downstream signaling in INS-1 cells and primary human pancreatic β-cells. We found that PDE1 and PDE4, but not PDE3 or PDE8, regulate resting and glucose-stimulated cAMP levels in INS-1 cells (Fig 1, S1 Fig), and this correlates with the ability of PDE1- and PDE4-selective inhibitors to enhance GSIS in these cells (Fig 4B). However, 8MM-IBMX was the only subtype-selective inhibitor to enhance glucose + forskolin-stimulated Ca^2+^ transients (Fig 5E&F), suggesting that PDE1 may regulate GSIS by modulating Ca^2+^ dynamics. In contrast to INS-1 cells, we found that PDE3 and PDE4, but not PDE1 or PDE8, regulate resting cAMP levels in primary human β-cells (Fig 2, S2 Fig). Strikingly, inhibition of PDE1 and PDE4 activity significantly increased cAMP accumulation following stimulation with 16.7 mM glucose in human β-cells (Fig 3). Consistent with this, the PDE1-selective inhibitor 8MM-IBMX had the greatest effect on GSIS in human islets (Fig 4C). Finally, we found that PDE1 inhibition reduced palmitate-induced caspase-3/7 activation (Fig 6B) and induced CREB phosphorylation (Fig 6C) in INS-1 cells. Taken together, our results suggest that PDE1-mediated regulation of cAMP levels is important for regulation of GSIS and pancreatic β-cell survival.

### PDE-mediated regulation of resting and glucose-stimulated cAMP levels in INS-1 cells and human pancreatic β-cells

We elected to study the role of PDE1, PDE3, PDE4, and PDE8 in PDE-mediated regulation of cAMP levels because they are among the most highly-expressed PDEs in pancreatic β-cell lines and islets from rat, mouse, and human (21, 28, 30, 41). Subtype-selective inhibitors and siRNA-mediated knockdown have revealed an important role for PDE1, PDE3, PDE4, and PDE8 in regulation of resting cAMP levels and GSIS (21-23). However, individual studies have focused on one or two PDE subtypes and have not performed a direct comparison of PDE1, PDE3, PDE4, and PDE8. The few studies that have examined PDE1, PDE3, PDE4, and PDE8 differed substantially in approach from the current study. For example, subtype-selective inhibitors of PDE1 and PDE4, but not PDE3, elevated cAMP content in homogenates of the mouse insulinoma β-cell line βTC3, suggesting that PDE1 and PDE4 are required (21). However, studying the effect of PDE inhibitors on cAMP dynamics in living pancreatic β-cells in real-time, as we performed in this study, is more likely to reflect cAMP dynamics in vivo. A systematic comparison of PDE1, PDE3, PDE4, and PDE8 was performed in the mouse insulinoma cell line MIN6 and primary mouse pancreatic β-cells using a plasma membrane-localized fluorescent cAMP sensor and TIRF microscopy (22). The authors concluded that PDE3 is the primary regulator of resting cAMP levels. However, PDE3 is commonly associated with cellular membranes, especially the plasma membrane (54, 55). PDE1 and PDE4, on the other hand, are associated with cellular sub-compartments, including the ER (56). Therefore, an approach that detects mainly sub-plasmalemmal cAMP may underestimate the contributions of PDE1 and PDE4. In the current study, we used a robust fluorescent cAMP sensor localized to the cytosol, Epac-S^H187^ (34), and found that under resting conditions, PDEs regulate resting cAMP levels in INS-1 cells with a rank order of PDE1 > PDE4 > PDE3 = PDE8 (Fig 1A&B, S1 Fig). In contrast, we found that in human pancreatic β-cells, PDE3 and PDE4, but not PDE1 or PDE8, regulate resting cAMP levels (Fig 2, S2 Fig). Two earlier studies have examined regulation of resting cAMP levels by PDE1, PDE3, and PDE4 in human islets (23, 41). However, the authors measured cAMP in cell lysates obtained from intact islets, which included contributions from α- and δ-cells, in addition to β-cells (57). Therefore, the present study, which specifically examines PDEs in isolated human pancreatic β-cells, is much less likely to be influenced by contributions from other cell types resident within the pancreatic islet.

While PDE-mediated regulation of resting cAMP levels has been extensively studied, regulation of cAMP levels by PDE1, PDE3, and PDE4 in the presence of glucose remains largely unexplored. This data is significant because it would be directly comparable with the effect of subtype-selective PDE inhibitors on GSIS. In this study, we found that stimulatory concentrations of glucose altered the PDE profile in INS-1 cells (Fig 1E) and human β-cells (Fig 3E). The IBMX-stimulated rise in cAMP levels was significantly greater in the absence of glucose than in the presence of 18 mM glucose in INS-1 cells (Fig 1E); however, this was not the case for the subtype-selective inhibitors. In fact, the PDE1-selective inhibitor 8MM-IBMX and PDE3-selective inhibitor cilostamide trended toward a greater effect in the presence of glucose. PDE activity is governed by the local cAMP concentration, which in turn is dependent on the presence of cAMP-generating AC subtypes within cAMP microdomains or compartments in the cell (58). PDE activity can also be regulated by protein kinase phosphorylation, such as PKA-mediated phosphorylation and upregulation of PDE3 and PDE4 activities (40). In addition, PDE1 activity is upregulated by Ca^2+^/Calmodulin (19, 20). The observation that the IBMX-stimulated rise in cAMP levels decreased in the presence of glucose, while the effect of subtype-selective inhibitors did not change, suggests that PDE1, PDE3 and PDE4 transition (Fig 1D) from regulating different cAMP compartments under resting conditions (subtype-selective inhibitors added up to IBMX) to regulating a similar cAMP microdomain during glucose stimulation (subtype-selective inhibitors are roughly equivalent to IBMX). Indeed, stimulus-induced changes in the components of cAMP microdomains, including PDEs, have been reported previously (56, 59). Glucose elevates Ca^2+^ and cAMP levels and stimulates PKA activity (8, 10, 12); therefore, glucose stimulation of PDE1 and PDE3 activities may be due to regulation by Ca^2+^/Calmodulin and PKA, respectively. In contrast with our findings in INS-1 cells, the IBMX-stimulated rise in cAMP levels was significantly elevated following glucose stimulation of human β-cells (Fig 3E). This was accompanied by a significant increase in the 8MM-IBMX- and rolipram-induced elevations in cAMP levels. PDE4 activity is upregulated by PKA- and CamKII-mediated phosphorylation (40, 60); therefore, it’s possible that glucose stimulates PKA- or CamKII-dependent phosphorylation of PDE4 in human β-cells. Our finding that PDE1 and PDE4, but not PDE3, activities are upregulated by glucose in human β-cells may also be explained in the context of the affinity of the different PDEs for cAMP. The resting cAMP concentration in mammalian cells is estimated to be near 1 µM (61, 62), close to the K_m_ for cAMP of PDE1C and PDE4 isoforms, below the K_m_ for cAMP of PDE1A and 1B, but much higher than that of all PDE3 isoforms, which have K_m_ values for cAMP of between 20 and 150 nM (63). Therefore, PDE3 activity is likely maximal under resting conditions, whereas the activities of PDE1 and PDE4 would be expected to increase in response to a glucose-stimulated elevation in cAMP levels.

### PDE-mediated regulation of Ca^2+^ dynamics and GSIS in INS-1 cells and human islets

We found that inhibition of PDE1 and PDE4 potentiated GSIS INS-1 cells (Fig 4B), while inhibition of PDE3 (Fig 4B) and PDE8 (S1 Fig) did not. Unfortunately, none of our experiments with human islets resulted in significant GSIS over basal secretion. However, in two donors (donors 4&6), IBMX increased insulin secretion over glucose alone, and in another donor (donor 3), inhibition of PDE1 significantly increased insulin secretion over glucose alone (Fig 4C). Inhibition of PDE1 trended toward potentiation of GSIS in two additional donors (donors 4&5). Our result in INS-1 cells is consistent with our finding that inhibition of PDE1 or PDE4 potentiates glucose-stimulated cAMP levels (Fig 1C&D). Other studies on PDE-regulation of GSIS demonstrate that PDE3 is involved in regulating GSIS (24, 26-28, 64). One exception to this is that the PDE1-selective inhibitor 8MM-IBMX was shown to potentiate GSIS in βTC3 cells (21), in agreement with our study. A direct comparison of GSIS regulation by PDE1, PDE3, PDE4, and PDE8 showed that knockdown of each subtype enhanced GSIS in INS-1 cells; however, there was no difference among them (30). Another study found that PDE1 is an important regulator of GSIS in MIN6 cells, but also reported that PDE3 and PDE8 regulate GSIS as well (22). In human islets PDE1, PDE3, and PDE4, together comprised the vast majority of total PDE activity, although other PDEs were detected by western blot (41). This is consistent with our conclusion that PDE1 inhibition can potentiate GSIS in human islets (Fig 4C).

The rise in cAMP levels stimulated by the pan PDE inhibitor IBMX, PDE1-selective inhibitor 8MM-IBMX and PDE4-selective inhibitor rolipram in INS-1 cells (Fig 1) and human pancreatic β-cells (Fig 2&3) likely resulted in activation of the cAMP effectors PKA and Epac (12); however, it is only in the presence of glucose that PDE inhibition has the ability to stimulate insulin secretion (65). PKA and Epac enhance insulin secretion at the site of exocytosis through phosphorylation of exocytotic substrates (13) and direct interactions with scaffolding proteins (66), SNARE proteins (67) and ion channels (15), respectively. PKA and Epac can also enhance insulin secretion through more distal actions on ER Ca^2+^ channels IP_3_R and RyR. PKA directly phosphorylates the IP_3_R to stimulate ER Ca^2+^ release (68-70), whereas Epac is reported to regulate RyR-mediated Ca^2+^ release through a Rap1/PLCε/CamKII signaling pathway (16, 71). Thus, we tested the possibility that PDE inhibition regulates [Ca^2+^]_in_ under various conditions. While we could not detect any change in cytosolic Ca^2+^ levels with the subtype-selective PDE inhibitors or IBMX in the absence or presence of 18 mM glucose, we found that the PDE1-selective inhibitor 8MM-IBMX potentiated the Ca^2+^ rise stimulated with 18mM glucose + the AC activator forskolin to a level comparable with IBMX (Fig 5E&F). We’ve previously reported that 25 μM forskolin, in the presence of IBMX, stimulates a robust increase in [cAMP] as assessed with the Epac-S^H187^ sensor in INS-1 cells (72). Thus, while the effect of IBMX and 8MM-IBMX on cAMP levels in INS-1 cells is readily detectable even under resting conditions (Fig 1A&B), they only significantly increase global Ca^2+^ levels over that stimulated by 18 mM glucose when transmembrane ACs are strongly stimulated. Interestingly, inhibition of neither PDE3 nor PDE4 potentiated the Ca^2+^ transient stimulated by 18 mM glucose + forskolin (Fig 5F). Together, our results suggest that inhibition of PDE1 could potentially enhance the efficacy of incretin hormones, such as GLP-1 or gastrointestinal inhibitory peptide, that activate transmembrane ACs through G-protein coupled receptors (73). In cardiomyocytes, cAMP microdomains containing PDEs, ACs, the cAMP effectors PKA and Epac, are anchored near target substrates, including ER Ca^2+^ release channels, by A-Kinase Anchoring Protein (AKAP) (74-76). Indeed, evidence of a cAMP microdomain containing AKAP150, PKA and AC8 has been demonstrated in pancreatic β-cells (77-79). Whether PDE1 is present, and the precise sub-cellular localization of this cAMP microdomain, remains to be determined.

### PDE inhibitors as a potential treatment for type 2 diabetes

As demonstrated by GLP-1 receptor stimulation, elevated cAMP levels not only enhance GSIS but can stimulate pancreatic β-cell proliferation (80) and cell survival (81). Indeed, the pan PDE inhibitor IBMX has the ability to rescue pancreatic β-cells from cell death (Fig 6B) induced by the unsaturated fatty acid palmitate (82). Palmitate exposure recapitulates key aspects of pancreatic β-cell dysfunction observed in diabetes, including disruption of glucose-stimulated cAMP oscillations (50, 51). We also showed that the PDE1-selective inhibitor 8MM-IBMX significantly reduces palmitate-induced caspase-3/7 activation in INS-1 cells, while the PDE3-selective inhibitor cilostamide and PDE4-selective inhibitor rolipram do not (Fig 6B). Consistent with our observation, PDE1A is strongly upregulated in INS-1 832/13 cells in response to 48 hr palmitate exposure (83).

Activation of the cAMP-response element binding protein (CREB) is associated with β-cell survival (7, 84). Furthermore, IBMX has been shown to induce CREB phosphorylation via a PKA-dependent mechanism in other cell types (85). In our study, we found that IBMX and 8MM-IBMX significantly stimulated CREB phosphorylation above resting levels, while cilostamide and rolipram did not (Fig 6C), suggesting that PDE1 inhibition may suppress pancreatic β-cell apoptosis via a CREB-dependent mechanism. Since GSIS and pancreatic β-cell mass are compromised in type 2 diabetics (2), it’s possible that PDE1 inhibitors could have clinical applications for treating this disease (86). A recent study demonstrated that IBMX, the PDE3 inhibitors cilostamide and milrinone, the PDE4 inhibitor rolipram, and the PDE4/10 inhibitor dipyridamole stimulated rat pancreatic β-cell replication, and that the latter reduces serum glucose levels in humans (6). However, the PDE1 inhibitor 8MM-IBMX did not significantly stimulate β-cell replication in this study, suggesting that PDE1 may regulate GSIS and pancreatic β-cell survival but not proliferation. It will be of interest to determine if specific PDE1 isoforms play distinct roles in the regulation of pancreatic β-cell function and survival.

## Acknowledgements

The authors thank Dr. Kees Jalink, The Netherlands Cancer Center, for the gift of the Epac-S^H187^ cAMP sensor.

## Author Contributions

Conceptualization: Pratt, Harvey, Hockerman

Investigation: Pratt, Harvey, Salyer, Tang

Formal Analysis: Pratt, Harvey, Hockerman

Funding Acquisition: Hockerman

Project Administration: Hockerman

Validation: Pratt, Harvey, Salyer

Writing-original draft: Pratt, Hockerman

## Supporting Information Captions

### S1 Table-Human Islet Donor Characteristics

**S1 Fig-PDE 8-selective inhibitors don’t potentiate cAMP levels or insulin secretion in INS-1 cells-** (A) Application of 100 nM PF4957325 on INS-1 resulted in no increase in cAMP over baseline. Subsequent application of 100 μM IBMX resulted in increased cAMP ranging from 7-49% over baseline (n= 14 cells) (B) PF4957325 does not potentiate glucose-stimulated insulin secretion in INS-1 cells. Stimulation of INS-1 cells with 18 mM glucose resulted in a significant increase in insulin secretion over basal secretion. Co-stimulation with 100 nM PF4957325 failed to result in a significant increase in secretion compared to 18 mM glucose alone(n=3). (C) PF04671536 does not increase intracellular cAMP in INS-1 cells. Application of increasing concentrations of PF04671536 failed to increase in cAMP over baseline. Application of 100 μM IBMX resulted in increases in cAMP ranging from 23-113% over baseline (n=11 cells).

**S2 Fig-The PDE8-selective inhibitor PF04671536 does not increase basal cAMP levels in human pancreatic β-cells.** Application of 100 nM PF04671536 does not result in any increase in cAMP over baseline. Application of 100 μM IBMX increased cAMP ranging from 40-101%. GLP-1 and clonidine were used to identify β-cells (n=4).

## References

1. Rorsman P, Eliasson L, Renstrom E, Gromada J, Barg S, Gopel S. The Cell Physiology of Biphasic Insulin Secretion. News Physiol Sci. 2000;15:72–7.

2. Weir GC, Bonner-Weir S. Five stages of evolving beta-cell dysfunction during progression to diabetes. Diabetes. 2004;53 Suppl 3:S16-21.

3. Ohta M, Nelson J, Nelson D, Meglasson MD, Erecinska M. Effect of Ca++ channel blockers on energy level and stimulated insulin secretion in isolated rat islets of Langerhans. J Pharmacol Exp Ther. 1993;264:35–40.

4. Johnson JD, Kuang S, Misler S, Polonsky KS. Ryanodine receptors in human pancreatic beta cells: localization and effects on insulin secretion. FASEB J. 2004;18:878–80.

5. Ramos LS, Zippin JH, Kamenetsky M, Buck J, Levin LR. Glucose and GLP-1 stimulate cAMP production via distinct adenylyl cyclases in INS-1E insulinoma cells. J Gen Physiol. 2008;132:329–38.

6. Zhao Z, Low YS, Armstrong NA, Ryu JH, Sun SA, Arvanites AC, et al. Repurposing cAMP-modulating medications to promote beta-cell replication. Mol Endocrinol. 2014;28:1682– 97.

7. Jhala US, Canettieri G, Screaton RA, Kulkarni RN, Krajewski S, Reed J, et al. cAMP promotes pancreatic beta-cell survival via CREB-mediated induction of IRS2. Genes Dev. 2003;17:1575–80.

8. Dyachok O, Idevall-Hagren O, Sagetorp J, Tian G, Wuttke A, Arrieumerlou C, et al. Glucose-induced cyclic AMP oscillations regulate pulsatile insulin secretion. Cell Metabol. 2008;8:26–37.

9. Kitaguchi T, Oya M, Wada Y, Tsuboi T, Miyawaki A. Extracellular calcium influx activates adenylate cyclase 1 and potentiates insulin secretion in MIN6 cells. Biochem J. 2013;450:365– 73.

10. Landa LR, Jr., Harbeck M, Kaihara K, Chepurny O, Kitiphongspattana K, Graf O, et al. Interplay of Ca2+ and cAMP signaling in the insulin-secreting MIN6 beta-cell line. J Biol Chem. 2005;280:31294–302.

11. Dyachok O, Isakov Y, Sagetorp J, Tengholm A. Oscillations of cyclic AMP in hormonestimulated insulin-secreting beta-cells. Nature. 2006;439:349–52.

12. Idevall-Hagren O, Barg S, Gylfe E, Tengholm A. cAMP mediators of pulsatile insulin secretion from glucose-stimulated single beta-cells. J Biol Chem. 2010;285:23007–18.

13. Song WJ, Seshadri M, Ashraf U, Mdluli T, Mondal P, Keil M, et al. Snapin mediates incretin action and augments glucose-dependent insulin secretion. Cell Metab. 2011;13:308–19.

14. Skelin M, Rupnik M. cAMP increases the sensitivity of exocytosis to Ca(2)+ primarily through protein kinase A in mouse pancreatic beta cells. Cell Calcium. 2011;49:89–99.

15. Kang G, Chepurny OG, Malester B, Rindler MJ, Rehmann H, Bos JL, et al. cAMP sensor Epac as a determinant of ATP-sensitive potassium channel activity in human pancreatic beta cells and rat INS-1 cells. J Physiol. 2006;573:595–609.

16. Dzhura I, Chepurny OG, Kelley GG, Leech CA, Roe MW, Dzhura E, et al. Epac2-dependent mobilization of intracellular Ca(2)+ by glucagon-like peptide-1 receptor agonist exendin-4 is disrupted in beta-cells of phospholipase C-epsilon knockout mice. J Physiol. 2010;588:4871–89.

17. Kang G, Joseph JW, Chepurny OG, Monaco M, Wheeler MB, Bos JL, et al. Epac-selective cAMP analog 8-pCPT-2’-O-Me-cAMP as a stimulus for Ca2+-induced Ca2+ release and exocytosis in pancreatic beta-cells. J Biol Chem. 2003;278:8279–85.

18. Tengholm A. Cyclic AMP dynamics in the pancreatic beta-cell. Ups J Med Sci. 2012;117:355–69.

19. Goraya TA, Cooper DM. Ca2+-calmodulin-dependent phosphodiesterase (PDE1): current perspectives. Cell Signal. 2005;17:789–97.

20. Goraya TA, Masada N, Ciruela A, Cooper DM. Sustained entry of Ca2+ is required to activate Ca^2+^-calmodulin-dependent phosphodiesterase 1A. J Biol Chem. 2004;279:40494–504.

21. Han P, Werber J, Surana M, Fleischer N, Michaeli T. The calcium/calmodulin-dependent phosphodiesterase PDE1C down-regulates glucose-induced insulin secretion. J Biol Chem 1999;274:22337–44.

22. Tian G, Sagetorp J, Xu Y, Shuai H, Degerman E, Tengholm A. Role of phosphodiesterases in the shaping of sub-plasma-membrane cAMP oscillations and pulsatile insulin secretion. J Cell Sci. 2012;125:5084–95.

23. Parker JC, VanVolkenburg MA, Ketchum RJ, Brayman KL, Andrews KM. Cyclic AMP phosphodiesterases of human and rat islets of Langerhans: contributions of types III and IV to the modulation of insulin secretion. Biochem Biophys Res Commun. 1995;217:916–23.

24. Harndahl L, Jing XJ, Ivarsson R, Degerman E, Ahren B, Manganiello VC, et al. Important role of phosphodiesterase 3B for the stimulatory action of cAMP on pancreatic beta-cell exocytosis and release of insulin. J Biol Chem. 2002;277:37446–55.

25. Walz HA, Wierup N, Vikman J, Manganiello VC, Degerman E, Eliasson L, et al. Beta-cell PDE3B regulates Ca2+-stimulated exocytosis of insulin. Cell Signal. 2007;19:1505–13.

26. Harndahl L, Wierup N, Enerback S, Mulder H, Manganiello VC, Sundler F, et al. Beta-cell-targeted overexpression of phosphodiesterase 3B in mice causes impaired insulin secretion, glucose intolerance, and deranged islet morphology. J Biol Chem. 2004;279:15214– 22.

27. Shafiee-Nick R, Pyne NJ, Furman BL. Effects of type-selective phosphodiesterase inhibitors on glucose-induced insulin secretion and islet phosphodiesterase activity. Brit J pharmacol. 1995;115:1486–92.

28. Dov A, Abramovitch E, Warwar N, Nesher R. Diminished phosphodiesterase-8B potentiates biphasic insulin response to glucose. Endocrinol. 2008;149:741–8.

29. Choi YH, Park S, Hockman S, Zmuda-Trzebiatowska E, Svennelid F, Haluzik M, et al. Alterations in regulation of energy homeostasis in cyclic nucleotide phosphodiesterase 3B-null mice. J Clin Invest. 2006;116:3240–51.

30. Waddleton D, Wu W, Feng Y, Thompson C, Wu M, Zhou YP, et al. Phosphodiesterase 3 and 4 comprise the major cAMP metabolizing enzymes responsible for insulin secretion in INS-1 (832/13) cells and rat islets. Biochem Pharmacol. 2008;76:884–93.

31. Zhao AZ, Bornfeldt KE, Beavo JA. Leptin inhibits insulin secretion by activation of phosphodiesterase 3B. J Clin Invest. 1998;102:869–73.

32. Zhao AZ, Zhao H, Teague J, Fujimoto W, Beavo JA. Attenuation of insulin secretion by insulin-like growth factor 1 is mediated through activation of phosphodiesterase 3B. Proc Natl Acad Sci U S A. 1997;94:3223–8.

33. Asfari M, Janjic D, Meda P, Li G, Halban PA, Wollheim CB. Establishment of 2-mercaptoethanol-dependent differentiated insulin-secreting cell lines. Endocrinol. 1992;130:167– 78.

34. Klarenbeek J, Goedhart J, van Batenburg A, Groenewald D, Jalink K. Fourth-Generation Epac-Based FRET Sensors for cAMP Feature Exceptional Brightness, Photostability and Dynamic Range: Characterization of Dedicated Sensors for FLIM, for Ratiometry and with High Affinity. PloS one. 2015;10:e0122513.

35. Hidaka H, Hayashi H, Kohri H, Kimura Y, Hosokawa T, Igawa T, et al. Selective inhibitor of platelet cyclic adenosine monophosphate phosphodiesterase, cilostamide, inhibits platelet aggregation. J Pharmacol Exp Ther. 1979;211:26–30.

36. Schwabe U, Miyake M, Ohga Y, Daly JW. 4-(3-Cyclopentyloxy-4-methoxyphenyl)-2-pyrrolidone (ZK 62711): a potent inhibitor of adenosine cyclic 3’,5’-monophosphate phosphodiesterases in homogenates and tissue slices from rat brain. Mol Pharmacol. 1976;12:900–10.

37. Ahn HS, Bercovici A, Boykow G, Bronnenkant A, Chackalamannil S, Chow J, et al. Potent tetracyclic guanine inhibitors of PDE1 and PDE5 cyclic guanosine monophosphate phosphodiesterases with oral antihypertensive activity. J Med Chem. 1997;40:2196–210.

38. Soderling SH, Beavo JA. Regulation of cAMP and cGMP signaling: new phosphodiesterases and new functions. Current opinion in cell biology. 2000;12:174–9.

39. Vang AG, Ben-Sasson SZ, Dong H, Kream B, DeNinno MP, Claffey MM, et al. PDE8 regulates rapid Teff cell adhesion and proliferation independent of ICER. PloS one. 2010;5(8):e12011.

40. Murthy KS, Zhou H, Makhlouf GM. PKA-dependent activation of PDE3A and PDE4 and inhibition of adenylyl cyclase V/VI in smooth muscle. Am J Physiol Cell Physiol. 2002;282:C508– 17.

41. Heimann E, Jones HA, Resjo S, Manganiello VC, Stenson L, Degerman E. Expression and regulation of cyclic nucleotide phosphodiesterases in human and rat pancreatic islets. PloS one. 2010;5:e14191.

42. DeNinno MP, Wright SW, Etienne JB, Olson TV, Rocke BN, Corbett JW, et al. Discovery of triazolopyrimidine-based PDE8B inhibitors: exceptionally ligand-efficient and lipophilic ligand-efficient compounds for the treatment of diabetes. Bioorg Med Chem Lett. 2012;22:5721–6.

43. Liu G, Jacobo SM, Hilliard N, Hockerman GH. Differential modulation of Cav1.2 and Cav1.3-mediated glucose-stimulated insulin secretion by cAMP in INS-1 cells: distinct roles for exchange protein directly activated by cAMP 2 (Epac2) and protein kinase A. J Pharmacol Exp Ther. 2006;318:152–60.

44. Truchan NA, Brar HK, Gallagher SJ, Neuman JC, Kimple ME. A single-islet microplate assay to measure mouse and human islet insulin secretion. Islets. 2015;7:e1076607.

45. Gromada J, Brock B, Schmitz O, Rorsman P. Glucagon-like peptide-1: regulation of insulin secretion and therapeutic potential. Basic Clin Pharmacol Toxicol. 2004;95:252–62.

46. Leech CA, Chepurny OG, Holz GG. Epac2-dependent rap1 activation and the control of islet insulin secretion by glucagon-like peptide-1. Vitam Horm. 2010;84:279–302.

47. Kong H, Jones PP, Koop A, Zhang L, Duff HJ, Chen SR. Caffeine induces Ca2+ release by reducing the threshold for luminal Ca2+ activation of the ryanodine receptor. Biochem J. 2008;414:441–52.

48. Wang Y, Jarrard RE, Pratt EP, Guerra ML, Salyer AE, Lange AM, et al. Uncoupling of Cav1.2 from Ca(2+)-induced Ca(2+) release and SK channel regulation in pancreatic beta-cells. Mol Endocrinol. 2014;28:458–76.

49. Garber AJ. Incretin effects on beta-cell function, replication, and mass: the human perspective. Diabetes Care. 2011;34 Suppl 2:S258-63.

50. Tian G, Sol ER, Xu Y, Shuai H, Tengholm A. Impaired cAMP Generation Contributes to Defective Glucose-Stimulated Insulin Secretion After Long-term Exposure to Palmitate. Diabetes. 2015;64:904–15.

51. Biden TJ, Boslem E, Chu KY, Sue N. Lipotoxic endoplasmic reticulum stress, beta cell failure, and type 2 diabetes mellitus. Trends Endocrinol Metab. 2014;25:389–98.

52. Hoppa MB, Collins S, Ramracheya R, Hodson L, Amisten S, Zhang Q, et al. Chronic palmitate exposure inhibits insulin secretion by dissociation of Ca(2+) channels from secretory granules. Cell Metab. 2009;10:455–65.

53. Brentnall M, Rodriguez-Menocal L, De Guevara RL, Cepero E, Boise LH. Caspase-9, caspase-3 and caspase-7 have distinct roles during intrinsic apoptosis. BMC Cell Biol. 2013;14:32.

54. Penmatsa H, Zhang W, Yarlagadda S, Li C, Conoley VG, Yue J, et al. Compartmentalized cyclic adenosine 3’,5’-monophosphate at the plasma membrane clusters PDE3A and cystic fibrosis transmembrane conductance regulator into microdomains. Mol Biol Cell. 2010;21:1097–110.

55. Shakur Y, Takeda K, Kenan Y, Yu ZX, Rena G, Brandt D, et al. Membrane localization of cyclic nucleotide phosphodiesterase 3 (PDE3). Two N-terminal domains are required for the efficient targeting to, and association of, PDE3 with endoplasmic reticulum. J Biol Chem. 2000;275:38749–61.

56. Carlisle Michel JJ, Dodge KL, Wong W, Mayer NC, Langeberg LK, Scott JD. PKA-phosphorylation of PDE4D3 facilitates recruitment of the mAKAP signalling complex. Biochem J. 2004;381:587–92.

57. Brissova M, Fowler MJ, Nicholson WE, Chu A, Hirshberg B, Harlan DM, et al. Assessment of human pancreatic islet architecture and composition by laser scanning confocal microscopy. J Histochem Cytochem. 2005;53:1087–97.

58. Baillie GS. Compartmentalized signalling: spatial regulation of cAMP by the action of compartmentalized phosphodiesterases. FEBS J. 2009;276:1790–9.

59. Richter W, Day P, Agrawal R, Bruss MD, Granier S, Wang YL, et al. Signaling from beta1- and beta2-adrenergic receptors is defined by differential interactions with PDE4. EMBO J. 2008;27:384–93.

60. Mika D, Richter W, Conti M. A CaMKII/PDE4D negative feedback regulates cAMP signaling. Proc Natl Acad Sci U S A. 2015;112:2023–8.

61. Borner S, Schwede F, Schlipp A, Berisha F, Calebiro D, Lohse MJ, et al. FRET measurements of intracellular cAMP concentrations and cAMP analog permeability in intact cells. Nat Protoc. 2011;6:427–38.

62. Koschinski A, Zaccolo M. Activation of PKA in cell requires higher concentration of cAMP than in vitro: implications for compartmentalization of cAMP signalling. Sci Rep. 2017;7:14090.

63. Omori K, Kotera J. Overview of PDEs and their regulation. Circ Res. 2007;100:309–27.

64. Walz HA, Harndahl L, Wierup N, Zmuda-Trzebiatowska E, Svennelid F, Manganiello VC, et al. Early and rapid development of insulin resistance, islet dysfunction and glucose intolerance after high-fat feeding in mice overexpressing phosphodiesterase 3B. J Endocrinol. 2006;189:629–41.

65. Komatsu M, Sato Y, Yamada S, Yamauchi K, Hashizume K, Aizawa T. Triggering of insulin release by a combination of cAMP signal and nutrients: an ATP-sensitive K+ channel-independent phenomenon. Diabetes. 2002;51 Suppl 1:S29-32.

66. Shibasaki T, Sunaga Y, Fujimoto K, Kashima Y, Seino S. Interaction of ATP sensor, cAMP sensor, Ca^2+^ sensor, and voltage-dependent Ca^2+^ channel in insulin granule exocytosis. J Biol Chem. 2004;279:7956–61.

67. Vikman J, Svensson H, Huang YC, Kang Y, Andersson SA, Gaisano HY, et al. Truncation of SNAP-25 reduces the stimulatory action of cAMP on rapid exocytosis in insulin-secreting cells. Am J Physiol Endocrinol Metab. 2009;297:E452–61.

68. Dyachok O, Tufveson G, Gylfe E. Ca^2+^-induced Ca^2+^ release by activation of inositol 1,4,5-trisphosphate receptors in primary pancreatic beta-cells. Cell Calcium. 2004;36:1–9.

69. Holz GG, Leech CA, Heller RS, Castonguay M, Habener JF. cAMP-dependent mobilization of intracellular Ca^2+^ stores by activation of ryanodine receptors in pancreatic beta-cells. A Ca^2+^ signaling system stimulated by the insulinotropic hormone glucagon-like peptide-1-(7-37). J Biol Chem. 1999;274:14147–56.

70. Soulsby MD, Wojcikiewicz RJ. The type III inositol 1,4,5-trisphosphate receptor is phosphorylated by cAMP-dependent protein kinase at three sites. Biochem J. 2005;392:493–7.

71. Dzhura I, Chepurny OG, Leech CA, Roe MW, Dzhura E, Xu X, et al. Phospholipase C-epsilon links Epac2 activation to the potentiation of glucose-stimulated insulin secretion from mouse islets of Langerhans. Islets. 2011;3:121–8.

72. Pratt EP, Salyer AE, Guerra ML, Hockerman GH. Ca^2+^ influx through L-type Ca^2+^ channels and Ca^2+^-induced Ca^2+^ release regulate cAMP accumulation and Epac1-dependent ERK 1/2 activation in INS-1 cells. Mol Cell Endocrinol. 2016;419:60–71.

73. Velasco M, Diaz-Garcia CM, Larque C, Hiriart M. Modulation of Ionic Channels and Insulin Secretion by Drugs and Hormones in Pancreatic Beta Cells. Mol Pharmacol. 2016;90:341–57.

74. Dodge KL, Khouangsathiene S, Kapiloff MS, Mouton R, Hill EV, Houslay MD, et al. mAKAP assembles a protein kinase A/PDE4 phosphodiesterase cAMP signaling module. EMBO J. 2001;20:1921–30.

75. Dodge-Kafka KL, Soughayer J, Pare GC, Carlisle Michel JJ, Langeberg LK, Kapiloff MS, et al. The protein kinase A anchoring protein mAKAP coordinates two integrated cAMP effector pathways. Nature. 2005;437:574–8.

76. Nijholt IM, Dolga AM, Ostroveanu A, Luiten PG, Schmidt M, Eisel UL. Neuronal AKAP150 coordinates PKA and Epac-mediated PKB/Akt phosphorylation. Cell Signal. 2008;20:1715–24.

77. Hinke SA, Navedo MF, Ulman A, Whiting JL, Nygren PJ, Tian G, et al. Anchored phosphatases modulate glucose homeostasis. EMBO J. 2012;31:3991–4004.

78. Lester LB, Langeberg LK, Scott JD. Anchoring of protein kinase A facilitates hormone-mediated insulin secretion. Proc Natl Acad Sci U S A. 1997;94:14942–7.

79. Willoughby D, Masada N, Wachten S, Pagano M, Halls ML, Everett KL, et al. AKAP79/150 interacts with AC8 and regulates Ca^2+^-dependent cAMP synthesis in pancreatic and neuronal systems. J Biol Chem. 2010;285:20328–42.

80. Rabinovitch A, Blondel B, Murray T, Mintz DH. Cyclic adenosine-3’,5’-monophosphate stimulates islet B cell replication in neonatal rat pancreatic monolayer cultures. J Clin Invest. 1980;66:1065–71.

81. Cunha DA, Ladriere L, Ortis F, Igoillo-Esteve M, Gurzov EN, Lupi R, et al. Glucagon-like peptide-1 agonists protect pancreatic beta-cells from lipotoxic endoplasmic reticulum stress through upregulation of BiP and JunB. Diabetes. 2009;58:2851–62.

82. Kwon G, Pappan KL, Marshall CA, Schaffer JE, McDaniel ML. cAMP Dose-dependently prevents palmitate-induced apoptosis by both protein kinase A- and cAMP-guanine nucleotide exchange factor-dependent pathways in beta-cells. J Biol Chem. 2004;279:8938–45.

83. Malmgren S, Spegel P, Danielsson AP, Nagorny CL, Andersson LE, Nitert MD, et al. Coordinate changes in histone modifications, mRNA levels, and metabolite profiles in clonal INS-1 832/13 beta-cells accompany functional adaptations to lipotoxicity. J Biol Chem. 2013;288:11973–87.

84. Costes S, Broca C, Bertrand G, Lajoix AD, Bataille D, Bockaert J, et al. ERK1/2 control phosphorylation and protein level of cAMP-responsive element-binding protein: a key role in glucose-mediated pancreatic beta-cell survival. Diabetes. 2006;55:2220–30.

85. Liu H, Tang JR, Choi YH, Napolitano M, Hockman S, Taira M, et al. Importance of cAMP-response element-binding protein in regulation of expression of the murine cyclic nucleotide phosphodiesterase 3B (Pde3b) gene in differentiating 3T3-L1 preadipocytes. J Biol Chem. 2006;281:21096–113.

86. Butler AE, Janson J, Bonner-Weir S, Ritzel R, Rizza RA, Butler PC. Beta-cell deficit and increased beta-cell apoptosis in humans with type 2 diabetes. Diabetes. 2003;52:102–10.

